# High-Resolution Spatial Transcriptomic Atlas of Mouse Soleus Muscle: Unveiling Single Cell and Subcellular Heterogeneity in Health and Denervation

**DOI:** 10.1101/2024.02.26.582103

**Authors:** Jer-En Hsu, Lloyd Ruiz, Yongha Hwang, Steve D. Guzman, Chun-Seok Cho, Weiqiu Cheng, Yichen Si, Peter Macpherson, Angelo Anacleto, Qingyang Zhao, Xiaoya Zhao, Mitchell Schrank, Goo Jun, Hyun-Min Kang, Myungjin Kim, Susan V. Brooks, Jun Hee Lee

## Abstract

Skeletal muscle exhibits pronounced cellular and subcellular heterogeneity, but comprehensive spatial mapping has been constrained by cell/nuclei dissociation-based methods that lose tissue architecture and by spatial platforms with insufficient resolution or limited transcriptome coverage. Here we present a high-resolution spatial transcriptomic atlas of mouse soleus muscle in longitudinal sections with unbiased whole-transcriptome coverage, enabling myofiber-resolved transcriptomes while preserving subcellular expression domains across the length of fibers. Combining histology-guided myofiber segmentation with unbiased grid-based mapping, we recover canonical fiber types and reveal widespread hybrid myofiber states in situ, including type IIb-associated signatures that are rare in soleus muscle and evident only when intramyofiber heterogeneity is assessed. At subcellular scale, we delineate the neuromuscular junction (NMJ) as a multi-compartment niche comprising postsynaptic myonuclei and spatially distinct peri-synaptic and myelinating Schwann cell-associated regions, each with characteristic gene programs. Applying this framework to denervation (3 and 7 days) identifies robust fiber-type-specific stress responses, coordinated remodeling of macrophage and fibroblast transcriptomes, and marked intramyofiber heterogeneity, including spatially nonuniform activation of damage-response genes along individual myofibers, with distinct transcriptional domains proximal and distal to the NMJ and associated degenerative histological features. Together, this atlas provides a high-resolution reference for muscle biology and clarifies how denervation reshapes myofiber, synaptic, and stromal-immune programs across cells and within cells in intact tissue.

## Introduction

Skeletal muscle plays a vital role in myriad physiological processes, including movement, posture maintenance, and metabolic regulation [1]. This complex tissue, characterized by its ability to convert chemical energy into mechanical force, is essential not only for physical activity but also for overall health and well-being. The architecture of skeletal muscle is remarkably intricate, featuring a variety of myofiber types, each with distinct functional and metabolic properties, and is interspersed with non-myocyte cells such as satellite cells, immune cells, fibroblasts, and endothelial cells. These diverse elements contribute to the muscle’s cellular heterogeneity, with each type playing a specific role in muscle physiology and response to stimuli [2–4].

Skeletal muscle exhibits a profound dependency on neural inputs; innervation is crucial for maintaining muscle tone and function [5]. Denervation, the loss of neural input, leads to rapid atrophy and functional impairment, a process that is not only relevant to traumatic injuries and in various neuromuscular diseases but also plays a significant role in the aging process [6]. This dynamic interplay between muscle and nerve highlights the importance of understanding muscle biology in a holistic manner. The molecular makeup of skeletal muscles, which includes a diverse array of proteins, enzymes, and signaling molecules, further underpins their dynamic adaptability to various physical demands and stressors, as well as their response to changes in innervation status. Thus, a comprehensive understanding of the detailed structure and function of skeletal muscle, encompassing its heterogeneity and complex interactions with neural inputs, is pivotal. Such knowledge is essential not only for advancing our understanding of muscle physiology but also for developing targeted therapies for muscle-related diseases and addressing the challenges of muscle deterioration with aging.

Single-cell transcriptomic technologies have brought significant insights into cellular heterogeneity across various biological systems, yet their application in skeletal muscle research is challenged by the unique nature of muscle tissue. The elongated and multinucleate structure of myofibers is not amenable to standard fluidics or microwell-based cell sorting techniques, which are designed for smaller, mononucleate cells. In response to these challenges, single-nucleus RNA sequencing (snRNA-seq) has emerged as an alternative, focusing on the analysis of individual nuclei instead of whole cells [4]. While snRNA-seq was successfully able to reveal transcriptome heterogeneity in muscle cells [7–11], it falls short in providing a complete picture of the transcriptomic activity of muscle fibers and overall tissues. Primarily, it captures only the nuclear transcriptome, potentially missing critical information contained within the cytoplasm. This exclusion is particularly significant in muscle fibers, where key elements of the cellular machinery and signaling pathways are located in the cytoplasm. More importantly, snRNA-seq provides no information on the spatial organization of these nuclei within different regions of the muscle fibers, an aspect crucial to understanding the complex interactions and functionality of skeletal muscle.

The limitations of single-cell and single-nucleus approaches have led researchers to explore spatial transcriptomics, a technique that preserves the spatial context of gene expression within tissues. Spatial transcriptomics provides a two-dimensional map of transcriptomic data, allowing researchers to visualize gene expression patterns in relation to the tissue architecture [12–14]. However, this method also faces significant challenges when applied to skeletal muscle research. One major limitation is the resolution; current spatial transcriptomics techniques, such as the 10X Visium platform, offer a resolution that is often too coarse to distinguish individual cells or subcellular structures, such as the intricate arrangement of muscle fibers and neuromuscular junctions, as in recently published studies [15–22]. This limitation means that the heterogeneity and complex interactions within skeletal muscle tissue can be obscured, with the resulting data representing a mixture of signals from multiple cell types. Furthermore, these data were captured from horizontal tissue sections, making it even more difficult to resolve cell types at specialized regions, such as the neuromuscular junctions. Consequently, while spatial transcriptomics adds valuable context to transcriptomic analysis, its current resolution constraints limit its ability to provide the detailed insights necessary for a complete understanding of the nuanced cellular and molecular dynamics of skeletal muscle.

To overcome the limitations of traditional single-cell and spatial transcriptomics in skeletal muscle research, we employed Seq-Scope, a cutting-edge method that dramatically enhances resolution and analytical depth [12–14, 23, 24]. Seq-Scope, characterized by its ability to provide ultra-high-resolution transcriptomic data, allows for the precise mapping of gene expression at the subcellular level. This advanced technology enabled us to conduct a comprehensive profiling of both normal skeletal muscle and its response to denervation along the longitudinal axis of muscle fibers. Our findings, with unprecedented detail, reveal not only the intricate structure of neuromuscular junctions (NMJs) but also previously underappreciated heterogeneity in the individual muscle fiber responses to denervation.

Furthermore, our high-resolution data illuminate the complex interactions between myofibers and non-myocyte cells, both of which undergo substantial changes in their transcriptome phenotypes during denervation responses. This study thus represents a significant advancement in the field of muscle biology, providing a foundation for future research into the molecular mechanisms of muscle function and disease.

## Results

### Overview of the Experiments and Dataset

Skeletal muscle poses unique challenges for single-cell and spatial transcriptomic analyses due to the extreme length and multinucleated architecture of myofibers [4, 7–11]. Among skeletal muscles, the mouse soleus represents an especially suitable model for spatial transcriptomic interrogation because of its mixed composition of type I and type II fibers, which exhibit distinct metabolic and functional properties [25]. Type I fibers are enriched in mitochondria and optimized for sustained, oxidative activity, whereas type II fibers encompass a spectrum of glycolytic and oxidative phenotypes adapted for rapid force generation [3]. In addition, the mouse soleus muscle is well characterized histologically and displays a relatively simple and ordered architecture, with individual fibers extending over more than 70% of the muscle length and aligned parallel to the longitudinal axis. Neuromuscular junctions are positioned at stereotyped locations along each fiber and collectively form a narrow transverse band across the muscle. These anatomical features make the mouse soleus muscle well suited for comprehensive spatial mapping of myofiber and synaptic transcriptomes.

Using the HiSeq implementation of Seq-Scope, which provides a continuous imaging area of approximately 1.5 mm × 60 mm [26], we profiled mouse soleus muscles sectioned along the longitudinal axis. To characterize denervation-induced remodeling, we analyzed muscles collected at 3 and 7 days following surgical denervation. In each animal, the left soleus muscle was denervated through sciatic nerve transection, while the contralateral right muscle served as an innervated control (Fig. 1A). In total, 32 longitudinal sections from 11 animals were analyzed, including untreated controls (C0; 3 mice, 13 sections), innervated controls at 3 days (C3; 1 mouse, 2 sections) and 7 days (C7; 1 mouse, 2 sections), as well as denervated muscles at 3 days (D3; 2 mice, 6 sections) and 7 days (D7; 4 mice, 9 sections) (Fig. 1B, 1C).

**Fig 1.**
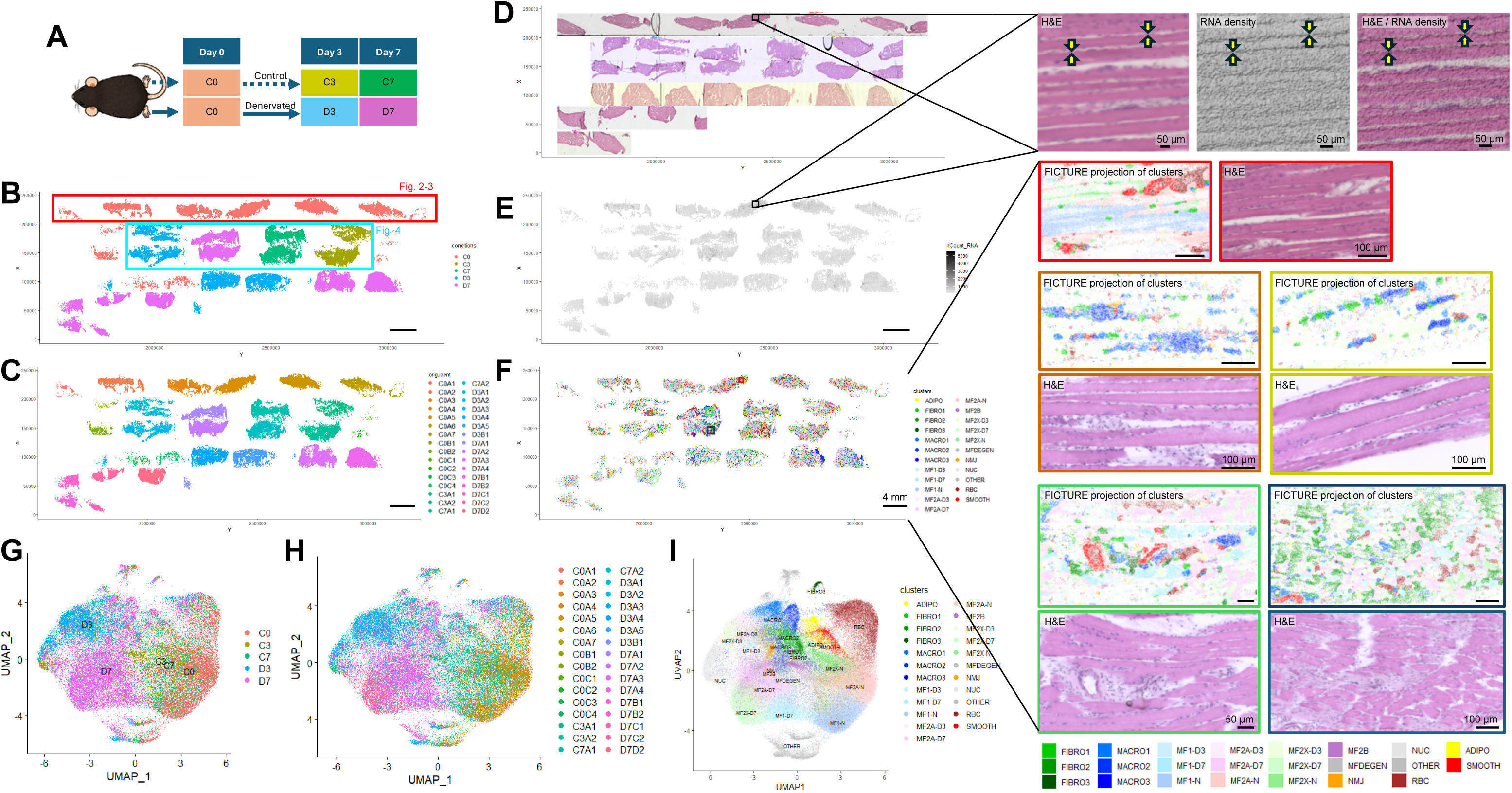
High-resolution spatial transcriptomic analysis of mouse soleus muscle in healthy and denervated conditions. (A) Schematic of the muscle denervation experiment. In each mouse, the left hindlimb was denervated, while the right hindlimb served as the control. Tissues were collected at baseline and at 3 and 7 days after denervation. (B-F) Spatial maps colored by experimental condition (B), individual section identity (C), hematoxylin and eosin (H&E) histology (D), captured RNA density (E), and transcriptome clustering results (F). Individual section identities in (C) are encoded by four letters, with the first two letters indicating experimental condition as in (A) and the third and fourth letters denoting animal and section identity, respectively. Boxed regions in (D-F) are shown at higher magnification on the right. Colored boxes in (F) correspond to magnified views of the same color, generated using pixel-level FICTURE projection of clusters. (G-I) UMAP embeddings colored by experimental condition (G), individual section identity (H), and transcriptome clustering results (I), corresponding to (B), (C), and (F), respectively.

Seq-Scope enables simultaneous acquisition of hematoxylin and eosin stained images and spatially resolved RNA capture from the same tissue section (Fig. 1D, 1E). Across all samples, we detected a total of 76.6 million unique reads mapped to the transcriptome. Spatial distributions of detected transcripts revealed a characteristic pattern, with lower transcript density in the central regions of myofibers and markedly higher density at the fiber periphery (Fig. 1D, 1E; magnified views). This pattern is consistent with muscle ultrastructure, in which the central fiber core is densely packed with myofibrils, whereas nuclei and most transcriptionally active cytoplasmic compartments are concentrated near the sarcolemma.

For downstream analysis, the spatial transcriptome was segmented into a grid of hexagonal bins with 14 µm sides. After excluding mitochondrial genes and hypothetical gene models and applying a minimum threshold of 100 detected gene features per bin to restrict analysis to tissue-covered regions (Fig. S1A), we obtained 121,191 hexagonal units. Each unit contained a median of 248 detected transcripts, with a mean ± SD of 311 ± 200 transcripts (Fig. S1B). These spatial units were subjected to unsupervised multidimensional clustering to identify major cellular and tissue compartments (Fig. 1F-1I).

Clustering analysis demonstrated clear separation between control tissues (C0, C3, and C7) and denervated tissues at 3 and 7 days (Fig. 1G). Importantly, biological and technical replicates within each experimental group showed strong integration, indicating minimal batch effects across experiments (Fig. 1H). These results confirm that the dataset robustly captures biologically meaningful transcriptional changes associated with denervation while maintaining high technical reproducibility.

Annotation of cluster identities revealed major myofiber types present in the soleus muscle, including type I, type IIa, and type IIx fibers, labeled as MF1, MF2A, and MF2X, respectively. In addition, multiple non-myocyte populations were identified, including macrophages, fibroblasts, smooth muscle cells, and adipocytes, each defined by canonical marker gene expression (Fig. 1I, Fig. S1C). Spatial projection of these clusters using FICTURE, a segmentation-free approach that infers transcriptomic patterns at submicrometer resolution [27], recapitulated the microscopic organization of muscle tissue observed by histology. This analysis accurately delineated myofiber boundaries and highlighted localized regions of immune infiltration, fibroblast expansion, and vascular structures (Fig. 1F, magnified panels).

Although non-myocyte populations were consistently detected across experimental conditions (Fig. S1D), tissue sections (Fig. S1E), and technical batches (Fig. S1F), myofiber clusters exhibited strong condition specificity. Distinct myofiber transcriptomic states were observed in control muscles (MF-N), 3-day denervated muscles (MF-D3), and 7-day denervated muscles (MF-D7) (Fig. S1D-S1F), indicating substantial denervation-induced alteration of myofiber gene expression. Application of batch correction and data integration methods successfully unified datasets across all conditions, recovering the canonical myofiber types (MF1, MF2A, and MF2X) and major non-myocyte populations covering all samples (Fig. S2). However, this integration also attenuated biologically meaningful condition-specific differences and eliminated populations unique to individual experimental states, underscoring the importance of complementary condition-aware analyses for capturing denervation-specific transcriptional programs.

### Single-Myofiber Transcriptome Analysis of Normal Muscle

From high-quality longitudinal sections of normal soleus muscle (Fig. 1B, red box), individual myofiber boundaries were delineated using H&E-guided segmentation (Fig. 2A) to enable single-myofiber transcriptome analysis. After applying a minimum cutoff of 300 detected gene features, 1,355 myofiber transcriptomes were obtained. Each myofiber revealed a median of 2,732 unique transcripts after excluding mitochondrial genes, hemoglobins, and hypothetical gene models (Fig. 2B, 2C). Variability in transcript counts reflected differences in the extent of longitudinal fiber representation within individual sections.

**Fig. 2.**
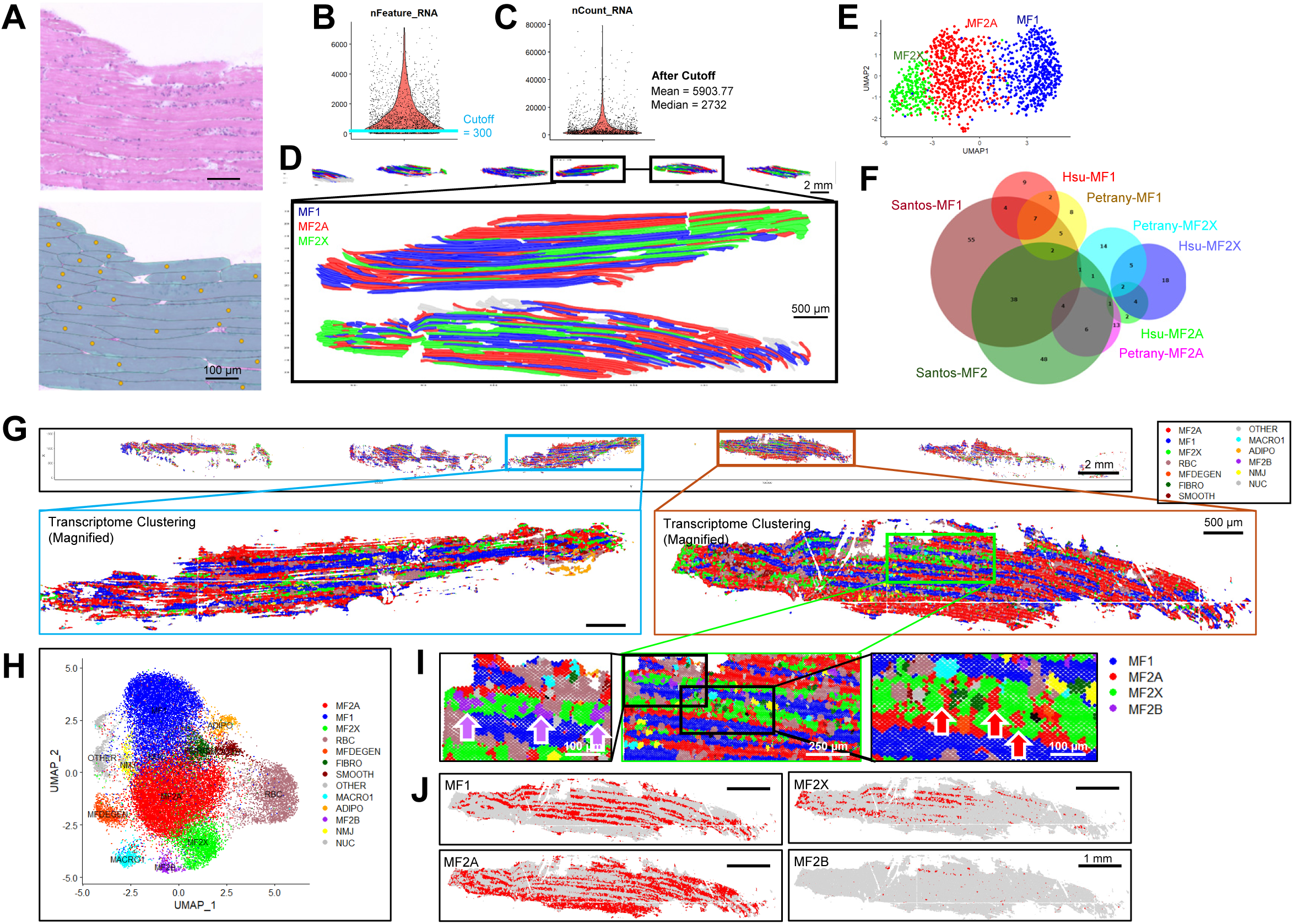
Single-myofiber transcriptome analysis of normal muscle. (A) Example of myofiber segmentation based on an H&E-stained image (top), with segmented fibers shown below. All transcriptomic data within each segmented region were aggregated into a single data point (yellow circle) for individual myofiber analysis. (B) Distribution of the number of gene features per segmented fiber (nFeature_RNA). A cutoff of 300 gene features was applied to retain high-quality transcriptomes. (C) Distribution of the number of unique transcripts per segmented fiber (nCount_RNA) after applying the 300 gene feature cutoff. (D) Projection of segmented myofiber transcriptome clusters from (E) into histological space based on the segmented regions. The upper panel shows the full analysis area, and the lower panels show magnified views of the fourth and fifth sections. (E) UMAP representation of the segmented myofiber transcriptome, revealing three major fiber types (MF1, MF2A, MF2X) by multidimensional clustering. (F) Comparison of marker gene sets for each myofiber type identified in the segmentation-based analysis with those reported in two prior single-nucleus RNA sequencing studies [7, 8], using a Venn diagram generated with DeepVenn [54]. (G) Spatial plot of a uniform hexagonal grid with 14 µm sides, with cell types projected from the clustering results in (H). Boxed regions are shown at higher magnification. (H) UMAP embedding of hexagonal grid-based transcriptome clustering colored by identified cell types. (I) Magnified regions from (G) highlighting hybrid fibers. (J) Spatial projection of individual myofiber type distributions, shown in red.

Unsupervised clustering resolved three major fiber type groups corresponding to MF1, MF2A, and MF2X (Fig. 2D, 2E). These groups were defined by expected fiber type-associated markers (Fig. S3A-S3C), including myosin heavy chain isoforms (Fig. S3D, first row), tropomyosins and troponins (Fig. S3D, second row), and ER calcium pumps and metabolic enzymes (Fig. S3D, third row). Fiber type-specific marker sets were largely consistent with prior snRNA-seq analyses of mouse soleus muscle [7, 8], while also revealing additional enriched genes not previously detected (Fig. 2F, Fig. S3E).

Because fiber identities were assigned in spatial context, each fiber type could be mapped back onto histological space, enabling direct visualization of their anatomical distribution (Fig. 2D). Inclusion of mitochondrial and hypothetical genes increased overall transcript counts but did not alter fiber type assignments (Fig. S3F). Notably, MF2X fibers expressed high levels of mitochondria-encoded transcripts despite their glycolytic role in soleus muscle (Fig. S3G, S3H)

### Subcellular Anatomy of the Hybrid Myofiber Transcriptome

To assess intra-myofiber transcriptomic heterogeneity, we analyzed spatial transcriptomes using 14 µm-sided hexagonal grids and focused this analysis on normal muscle tissue (Fig. 2G-2J, Fig. S4). Hexagonal clustering recapitulated the major myofiber types (Fig. 2H) and their characteristic markers (Fig. S4A), including myosin heavy chain isoforms (Fig. S4B), consistent with segmentation-based analysis (Fig. S4C) and prior snRNA-seq studies (Fig. S4D).

Despite this overall agreement, hexagonal clustering additionally detected small clusters with type IIb-associated signatures marked by *Myh4*, which are rare in the oxidative mouse soleus (Fig. 2H, Fig. S4A, S4B). To investigate this discrepancy, we examined the spatial distribution of hexagon-assigned myofiber types. High-resolution spatial mapping using multiscale sliding windows [24, 28] revealed that, while the overall spatial pattern of myofiber types resembled that obtained from segmentation-based analysis, hexagonal data uncovered transcriptomic heterogeneity within individual myofibers (Fig. 2G).

Specifically, distinct myofiber-type transcriptomes were detected within the same myofiber area, consistent with previously described hybrid myofibers [7, 8]. We observed mixtures of MF2A and MF2X, as well as MF2X and MF2B, within individual fibers (Fig. 2I). As expected given the rarity of MF2B fibers in mouse soleus, MF2B signatures were detected only in combination with MF2X within individual myofibers and were therefore not resolved as a distinct cluster by segmentation-based analysis, leading to their classification as MF2X (Fig. 2D, 2E).

In addition, we detected a small subset of hexagonal clusters characterized by elevated expression of cathepsin B, designated as MFDEGEN (Fig. S4E). Similar transcriptional signatures have been previously associated with exercise-induced remodeling [29] or muscle degeneration [30]. In normal muscle, MFDEGEN-associated regions were sparsely distributed and did not exhibit a consistent spatial pattern along myofibers, suggesting that this signature reflects a localized transcriptional state rather than a distinct fiber type or organized anatomical compartment.

### Identification of Neuromuscular Junction (NMJ) Transcriptome

Beyond myofiber type-associated transcriptomes, hexagonal binning of the Seq-Scope dataset revealed additional transcriptomic clusters corresponding to subcellular domains. Among these, a prominent cluster was characterized by high expression of the nuclear noncoding RNAs *Malat1* and *Neat1*, defining a nuclear (NUC) subdomain (Fig. 2H; Fig. 3A, blue) [31]. Adjacent to the NUC cluster in low-dimensional embedding, we identified a distinct transcriptomic cluster spatially localized to neuromuscular junctions (NMJs) (Fig. 3A, red). This cluster was defined by enrichment of canonical postsynaptic genes, including *Prkar1a, Etv5, Ufsp1*, and *Chrna1* (Fig. 3B), consistent with its localization to postsynaptic myonuclei at the NMJ. Spatial projection showed that NMJ-associated transcriptomic regions aligned precisely with the stereotyped midline localization of NMJs in soleus muscle (Fig. 3C). At higher magnification, these regions overlapped with histological endplate structures corresponding to clusters of postsynaptic myonuclei (Fig. 3D). NMJ markers identified in the Seq-Scope dataset showed partial overlap with postsynaptic nuclear markers previously identified by snRNA-seq (Fig. 3E), indicating that the NMJ transcriptome identified by Seq-Scope is consistent with known postsynaptic programs but also reveals distinct features not captured by snRNA-seq.

**Fig. 3.**
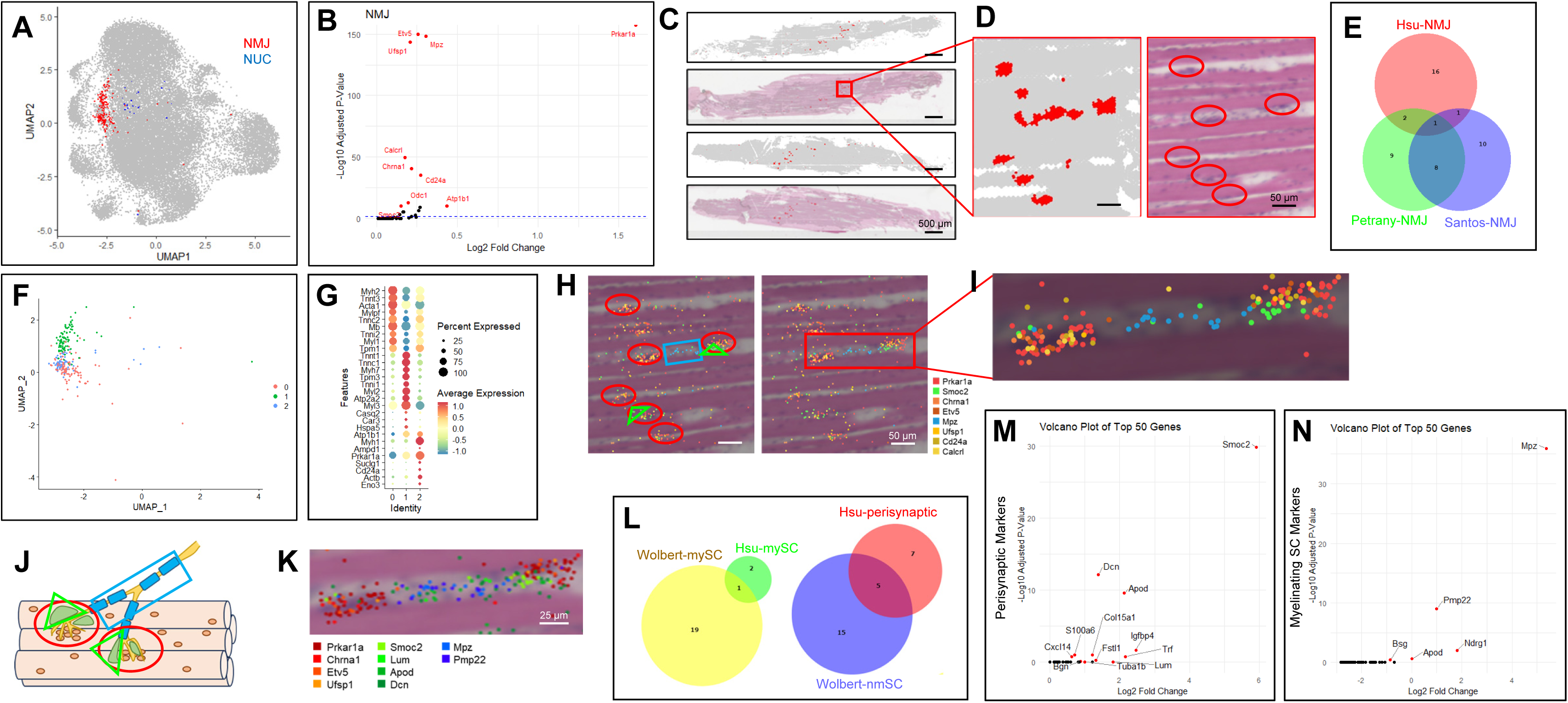
High-resolution spatial profiling of neuromuscular junction-associated transcriptomes. (A) Identification of neuromuscular junction (NMJ, red) and nuclear (NUC, blue) transcriptome clusters from the UMAP embedding of the hexagonal dataset. (B) Volcano plot showing genes enriched in the NMJ cluster. (C, D) Spatial plots of NMJ clusters shown in red in the first and third rows of (C), overlaid with histology in the second and fourth rows. The boxed region in (C) is magnified in (D, left) and shown with the corresponding H&E image (D, right). NMJ endplate structures in the histology are highlighted with red ovals. (E) Comparison of NMJ marker gene sets identified in the current hexagonal analysis with those from two previous single-nucleus RNA sequencing studies [7, 8], using a Venn diagram generated with DeepVenn [54]. (F, G) UMAP manifold of isolated NMJ transcriptomes (F), subclustered into three groups shown in different colors (0 = red, 1 = green, 2 = blue), corresponding to Type IIa, Type I, and Type IIx myofibers based on marker gene expression (G). These results suggest that postsynaptic NMJ nuclei retain their myofiber-type identities. (H-J) Spatial gene expression plots in the indicated colors, with gene detection represented as dots (H). Postsynaptic regions are indicated by red ovals, perisynaptic regions by green triangles, and myelinating Schwann cell regions by blue rectangles (H, left). The boxed region in (H, right) is magnified in (I). A schematic illustration of NMJ structure is shown in (J). (K-N) Identification of additional markers representing distinct NMJ-associated cell types. Markers enriched in the perisynaptic region (M) and myelinating Schwann cell region (N) are highlighted and compared with published marker lists (the top 20 significant genes) for non-myelinating and myelinating Schwann cells (L) [32]. Ten prominent markers were selected and projected onto the magnified NMJ spatial map (K), which displays architectural similarity to the canonical NMJ structure (J).

### Spatial Resolution of Postsynaptic and Schwann Cell-Associated NMJ Domains

Although the NMJ cluster was defined by postsynaptic gene expression, not all NMJ-associated transcriptomic regions overlapped with histological endplates. Several genes enriched in the NMJ cluster, including *Mpz* and *Smoc2*, are known markers of myelinating and non-myelinating Schwann cells, respectively [32]. Subclustering of the NMJ population did not separate these components and instead reflected fiber type-specific transcriptional signatures (Fig. 3F, 3G), indicating that NMJ-associated postsynaptic muscle transcriptomes retain fiber type-associated gene expression patterns.

We therefore examined the spatial distribution of individual NMJ marker genes with its raw resolution. This analysis revealed clear spatial separation between postsynaptic nuclear markers such as *Prkar1a, Etv5*, and *Ufsp1* and Schwann cell-associated markers *Mpz* and *Smoc2* (Fig. 3H, 3I). In contrast, acetylcholine receptor transcripts exhibited strong colocalization with postsynaptic nuclear regions marked by *Prkar1a* expression (Fig. S5A). These observations indicate that the NMJ-associated transcriptome comprises multiple spatially distinct domains that cannot be resolved by clustering alone but are separable through high-resolution spatial mapping.

### Distinct Transcriptomic Programs within the NMJ Microenvironment

Based on the spatial segregation of *Prkar1a*, *Smoc2*, and *Mpz*, we segmented three NMJ-associated regions representing postsynaptic myonuclei, a perisynaptic region, and a myelinating Schwann cell-associated region (Fig. 3H, 3J). Differential expression analysis between these compartments revealed distinct transcriptional programs (Fig. 3K-3N). The perisynaptic region exhibited high expression of *Lum, Apod, Dcn*, and *Smoc2* (Fig. 3M), closely matching non-myelinating Schwann cell transcriptomes described from brachial plexus and sciatic nerve tissues [32] (Fig. 3L, left). In contrast, the myelinating Schwann cell-associated region expressed genes involved in myelination, including *Mpz* [32], *Pmp22* [33], and *Ndrg1* [34] (Fig. 3N), showing strong concordance with myelinating Schwann cell transcriptomes derived from the same peripheral nerve dataset (Fig. 3L, right) [32].

Some of the genes we identified from the perisynaptic region, such as *ApoD* (Fig. 3M), were not identified in recently reported perisynaptic terminal Schwann cell transcriptome datasets [35, 36]. However, this expression pattern has been independently validated by RNAscope, which demonstrated high *ApoD* expression in terminal Schwann cells as well as in myelinating Schwann cells of intramuscular nerves [37]. In our analysis, *ApoD* emerged as a prominent marker of the perisynaptic region and was detected in both terminal and myelinating Schwann cell-associated regions (Fig. 3M, 3N). In addition, Smoc2 and Dcn, which were enriched in the perisynaptic region (Fig. 3M), are not only expressed in non-myelinating Schwann cells [32] but also in nerve-associated fibroblasts [38–40], indicating transcriptomic heterogeneity among cell populations adjacent to the NMJ.

Spatial visualization of compartment-specific markers revealed a geometric organization of postsynaptic nuclei, perisynaptic regions, and myelinating Schwann cell-associated domains that closely mirrors the known anatomical arrangement at the neuromuscular junction (Fig. 3J, Fig. S5B). Together, these results demonstrate that high-resolution Seq-Scope resolves the NMJ as a multi-compartment transcriptomic niche in intact skeletal muscle.

### Characterization of Non-Myocytes in Seq-Scope Dataset

High-resolution hexagonal analysis identified multiple non-myocyte transcriptomic populations within the soleus muscle, including macrophages, fibroblasts, adipocytes, smooth muscle cells, and erythrocytes (Fig. S6A-S6D). These populations exhibited distinct spatial localization patterns that aligned with tissue microanatomy. For example, smooth muscle transcriptomes were specifically localized around arterial structures (Fig. S6E, left panel), while adipocyte transcriptomes were confined to lipid-rich regions (Fig. S6E, right panel).

Seq-Scope enabled robust detection of cell populations that are difficult to capture using nucleus-based approaches. Macrophage transcriptomes, marked by *Cd74* and MHC components, were readily detected and spatially resolved across tissue sections (Fig. S6A-S6D), despite being largely absent from snRNA-seq datasets [7, 8]. Similarly, erythrocyte-associated transcripts, including hemoglobins, were uniquely identifiable due to the whole-transcriptome and spatial nature of Seq-Scope.

Fibroblast transcriptomes enriched for extracellular matrix genes, including *Dcn, Gsn*, and collagens, were broadly distributed in interstitial regions and showed close correspondence to fibro-adipogenic progenitor-like signatures described in snRNA-seq studies (Fig. S6F). Likewise, smooth muscle-specific gene expression, including *Acta2, Myl9, Tagln*, and *Mylk* (Fig. S6D), were robustly detected, while prior snRNA-seq datasets isolated different set of markers (Fig. S6G). In contrast, certain cell types reported in snRNA-seq datasets, such as satellite cells, myotendinous junction-associated nuclei, and tenocytes, were underrepresented in the Seq-Scope dataset, likely reflecting volumetric sampling limitations inherent to thin tissue sections.

Taken together, analyses of normal soleus muscle demonstrate that Seq-Scope captures myofiber type diversity, intra-fiber transcriptomic heterogeneity, neuromuscular junction subdomains, and multiple non-myocyte populations within intact tissue. This spatially resolved baseline provides a reference for examining how these transcriptional programs and tissue architectures are altered following denervation.

### Fiber type-Specific Responses to Denervation

To examine fiber type-specific responses to denervation, we performed another round of segmentation-based single myofiber analysis, using four tissues that produced high-quality histology and transcriptome data (Fig. 4A). After applying a gene feature cutoff of 300, we obtained 3908 segmented myofiber transcriptomes from control and denervated fibers at 3 days (1337 and 620, respectively) and 7 days (1127 and 824, respectively). After removing all mitochondrial, hypothetical and hemoglobin-encoding transcripts, the median and mean numbers of transcripts in each myofiber was 1738 and 3274 ± 5332 respectively (Fig. 4B). As observed from hexagonal analysis (Fig. 1), denervated fibers at 3 and 7 days exhibited transcriptomes substantially different from control fibers, while fibers from control muscles at 3 and 7 days integrated very well into each other (Fig. S7A), indicating that the experimental batch effect is minimal and the differences we observed could be explained largely by the biological effects of denervation. Some of the fibers in day 7 denervated fibers contained a fibrotic signature, likely due to effects from fibroblast transcriptome nearby the myofibers. As we focused on myofiber transcriptomes, the fibers displaying fibrotic signatures were excluded from the fiber type-specific analysis (FIBRO in Fig. 4B; NA in Fig. 4C). In all muscles, we were able to identify three major myofiber types, corresponding to MF1, MF2A and MF2X fibers (Fig. 4B), based on fiber-type markers isolated from normal muscle (Fig. S7B).

**Fig. 4.**
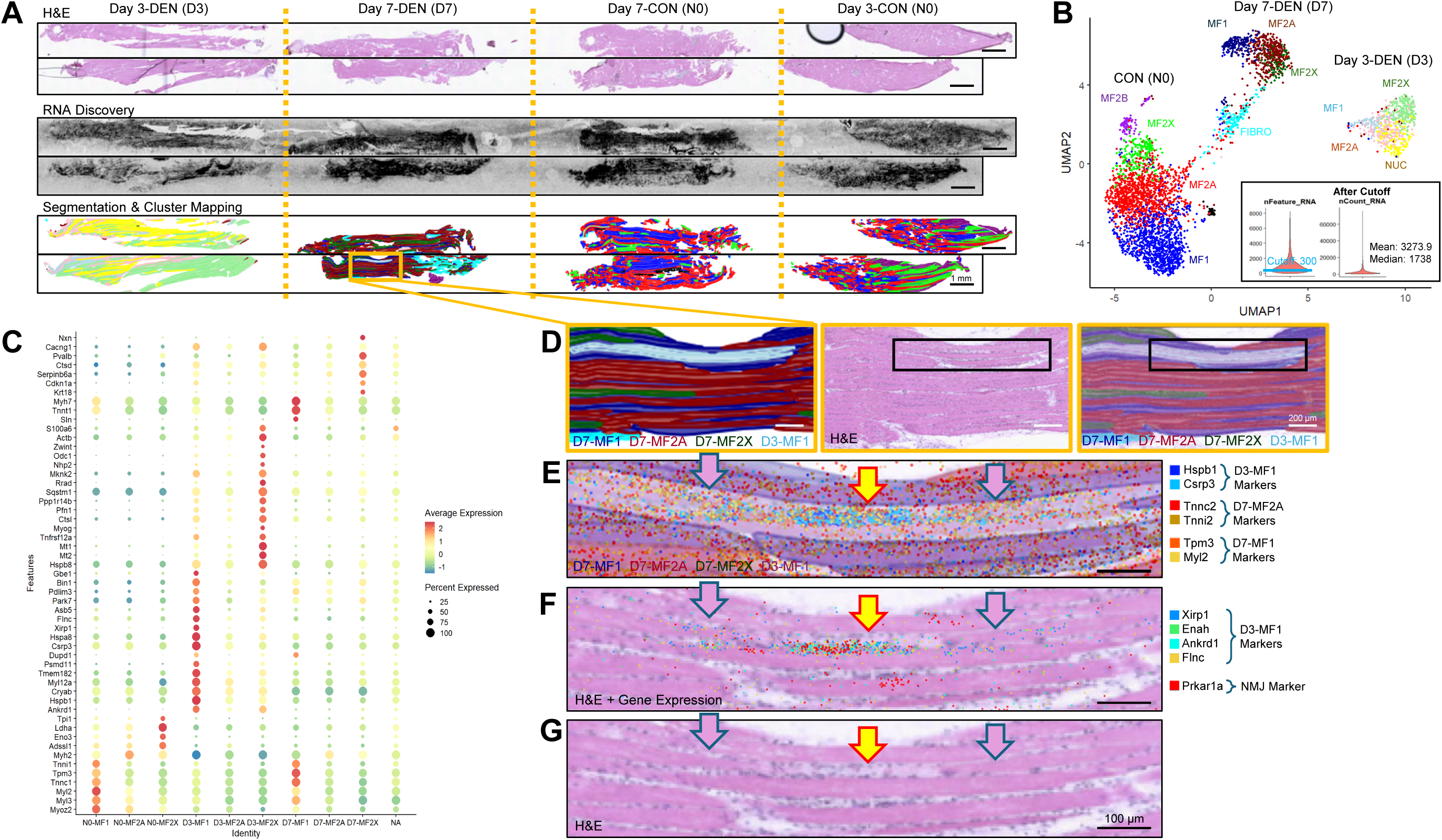
Heterogeneous transcriptome responses to denervation in single myofibers. (A) Longitudinal sections of mouse skeletal muscle from different denervation groups (DEN, denervated leg; CON, control leg), arranged as H&E histology (top), spatial RNA discovery plots (middle), and H&E-based segmented myofibers (bottom) projected with the fiber clustering colors shown in (B). (B) UMAP manifold of segmented myofiber transcriptomes, revealing heterogeneous denervation responses within three major fiber types identified by multidimensional clustering. Insets show the distribution of gene features per fiber (nFeature_RNA, left) and the distribution of unique transcript counts per fiber (nCount_RNA, right) after applying the gene feature cutoff of 300. (C) Visualization of cluster-specific top marker gene expression across myofiber clusters, highlighting fiber type-specific responses to denervation. (D-G) Identification of myofibers exhibiting intracellular heterogeneity in transcriptomic responses to denervation. Myofiber transcriptomes identified by segmentation of the H&E image (D, center) are projected into histological space (D, left) and overlaid onto the corresponding H&E image (D, right). Although the tissue was collected 7 days after denervation, a subset of myofibers exhibits a D3-MF1 (day 3 MF1-type) transcriptomic signature. The boxed region in (D) is magnified in (E-G). Selected denervation-induced genes and NMJ markers are plotted in the magnified regions (E, F), revealing close spatial proximity between degenerative responses and the NMJ area. Overlay with H&E histology shows that regions with high D3-MF1 signature (yellow arrow) display pale eosin staining, whereas regions exhibiting a D7-MF1 transcriptome (pink arrows) show normal myofiber staining (G).

Even though classical markers, myosin heavy chain, troponin and tropomyosin isoforms were relatively highly expressed in their corresponding myofiber types within the same section, allowing us to identify their original fiber types, their level was strongly diminished in day 3 denervated myofibers compared to control myofibers (Fig. S7B). In contrast, differential expression analyses identified a number of genes that were strongly upregulated in day 3 denervated fibers, but also explicitly upregulated in specific myofiber types, such as oxidative type I or glycolytic type IIx fibers (Fig. 4C, S7C). Markers enriched for day 3 denervated myofiber type I (D3-MF1) include several heat shock proteins (e.g. Hspb1, Hspa8), crystallins (*Cryab*) and muscle stress response proteins (e.g. *Csrp3, Ankrd1*). These were proteins previously shown to be upregulated upon denervation through proteomics [41]; however, our analyses show that their upregulation is highly specific to oxidative MF1 fibers (Fig. 4C, S7C, S7D). In addition to these genes, many markers of damaged myofibers (e.g. *Xirp1, Flnc, Enah* and *Otud1*; see references [7, 9, 42]) were also specifically upregulated in D3-MF1 (Fig. S7D). In contrast, day 3 denervated myofiber type IIx (D3-MF2X) exhibited stronger upregulation of different genes such as heat shock proteins (e.g. *Hspb8*) and metallothioneins (*Mt1, Mt2*) (Fig. S7E).

We also identified markers enriched for day 7 denervated fibers. Some classical type I myofiber markers, such as *Myh7* and *Tnnt1*, were upregulated in D7-MF1 (Fig. S7F), while other genes such as *Krt18*, *Cdkn1a* and *Ctsb* were upregulated in D7-MF2X (Fig. S7G). In addition to these, day 7 denervated fibers exhibited upregulation of various stress-responding genes such as autophagy controllers (Fig. S7H), atrophy inducers (Fig. S7I), a chromatin modulator (*Hdac3*) and oxygen metabolism genes (*Mb* and *Car3*; Fig. S7J), as well as DNA damage-responsive transcripts (*Cdkn1a, Gadd45a* and *Mdm2*; Fig. S7K), which show variable fiber type specificities in regulation.

### Subcellular Heterogeneity in Denervation Responses

Myofiber damage-response markers enriched in the D3-MF1 state, including *Xirp1, Enah*, and *Flnc* [42–44], have been reported previously in aged muscle and in myofibers undergoing dystrophic remodeling [7, 9], suggesting that their induction reflects acute structural or mechanical stress. Although the transcriptomic state defined as D3-MF1 was primarily observed in muscle tissue at 3 days after denervation, it was also detected in a subset of fibers at 7 days (Fig. 4D), indicating that the response is not exclusively specific to the early phase and can persist to the chronic phase. High-resolution spatial analysis revealed that these fibers often exhibited hybrid transcriptional states, with discrete intramyofiber regions expressing either D3-MF1 or D7-MF1 gene programs (Fig. 4E).

Importantly, these transcriptional domains were associated with distinct histological features. Regions expressing D3-MF1 markers showed pale and degenerative H&E staining patterns (Fig. 4E-4G, yellow arrows), whereas adjacent regions expressing D7-MF1 markers retained a more preserved myofiber morphology (Fig. 4E-4G, blue arrows). This spatial concordance between transcriptomic state and histological appearance indicates that denervation elicits localized, rather than uniform, remodeling responses along individual myofibers.

Notably, D3-MF1 damage-response transcripts were frequently enriched near the neuromuscular junction. While postsynaptic NMJ markers such as *Prkar1a* remained sharply confined to the endplate region, D3-MF1-associated transcripts extended beyond the synaptic core and radiated into adjacent segments of the same myofiber (Fig. 4F). This pattern suggests that early denervation-induced transcriptional responses are spatially organized around the NMJ and propagate into peri-synaptic regions, even in the absence of active neural input. Together, these findings demonstrate that denervation triggers pronounced subcellular heterogeneity within myofibers, linking localized structural degeneration, spatially restricted transcriptional programs, and the anatomical position of the NMJ.

### Denervation Preserves Postsynaptic NMJ Identity while Inducing Extrajunctional Acetylcholine Receptors

We next examined postsynaptic NMJ responses to denervation by focusing on the NMJ cluster identified in the hexagonal grid analysis covering the entire dataset (Fig. 5A; Fig. 1I). In the UMAP manifold, NMJ-associated transcriptomes were distributed across multiple subclusters, reflecting their strong similarity to the surrounding myofiber transcriptomes within the same tissue sections (Fig. 5A). Consistent with global myofiber responses (Fig. 4), NMJ-associated regions exhibited induction of stress-responsive genes, including heat shock proteins and crystallins, following denervation (Fig. S8A, S8B).

**Fig. 5.**
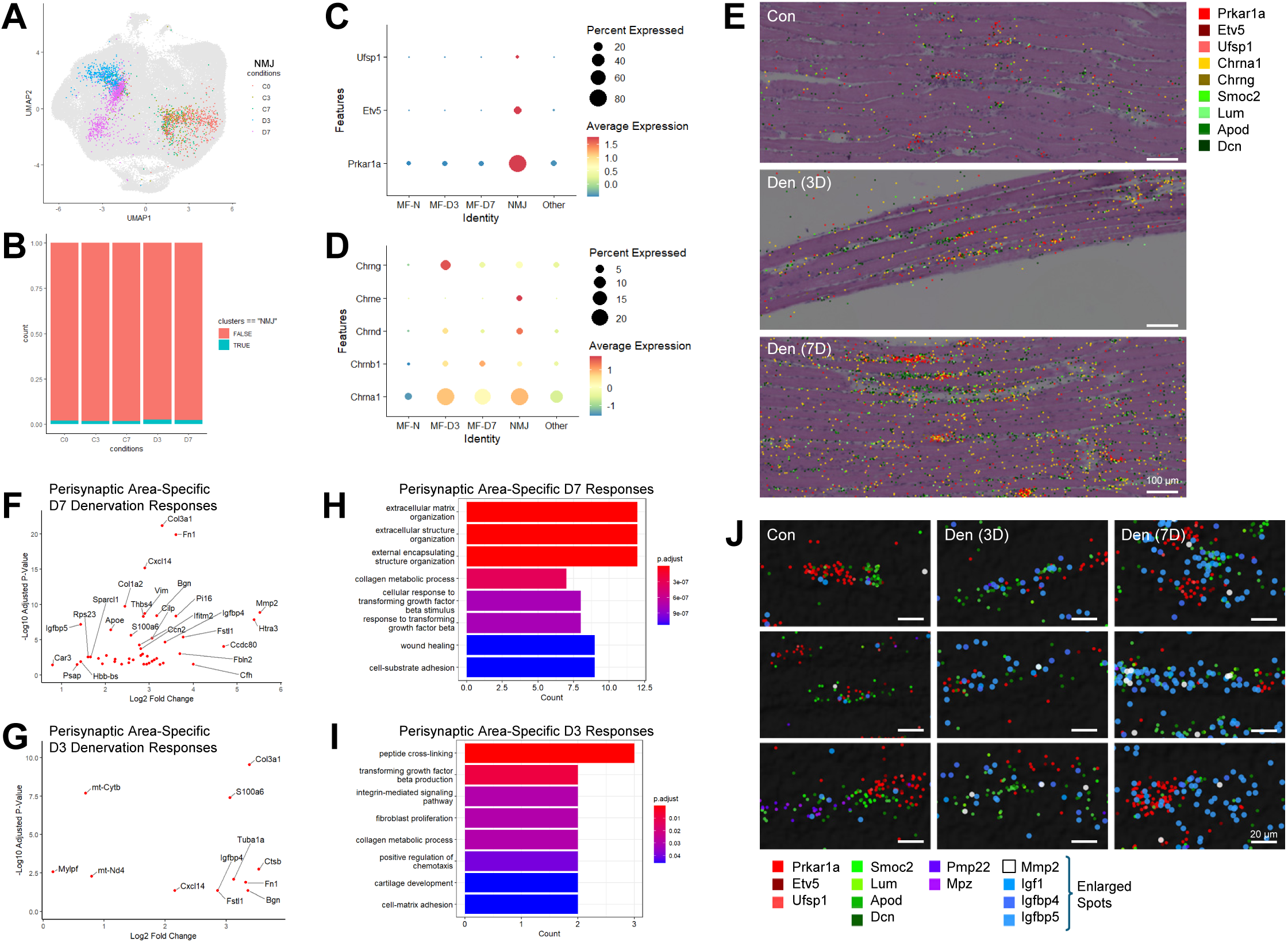
Denervation-induced changes in the neuromuscular junction region. (A) UMAP manifold from Fig. 1I highlighting the NMJ cluster, colored by experimental condition. All other clusters are shown in gray in the background. (B) Relative abundance of the NMJ cluster across experimental conditions. (C, D) Dot plots displaying differential expression of postsynaptic nuclei marker genes (C) and acetylcholine receptor isoforms (D) across myofiber types at different time points after denervation. (E) Spatial expression of postsynaptic nuclei markers (warm colors) and non-myelinating perisynaptic Schwann cell markers (greens) in control, 3-day denervated, and 7-day denervated myofibers, overlaid onto corresponding H&E images. (F-I) Perisynaptic area-specific transcriptional responses at 7 days (F, H) and 3 days (G, I) after denervation, identified by comparison with control perisynaptic regions. Genes that were also regulated in the postsynaptic area were excluded from the analysis. Results are shown as volcano plots (F, G) and gene ontology enrichment analyses (H, I). (J) Spatial expression of postsynaptic nuclei markers (reds), perisynaptic area markers (greens), myelinating Schwann cell area markers (purples), Mmp2 (white), and Igf and Igfbp genes (blues), indicating enhanced extracellular matrix remodeling and IGF pathway activity around the NMJ following denervation.

Despite these changes, the transcriptional identity of postsynaptic NMJs remained remarkably stable. Core postsynaptic markers, including *Prkar1a, Etv5*, and *Ufsp1*, maintained robust and spatially restricted expression at the endplate throughout the denervation time course (Fig. 5B, 5C; Fig. S8D). The overall abundance of NMJ-associated regions in the dataset was not substantially altered by denervation (Fig. 5B), indicating preservation of NMJ structural organization at the transcriptional level.

In contrast, acetylcholine receptor gene expression underwent marked remodeling, consistent with established models where neuronal activity suppresses acetylcholine receptor transcription in extrasynaptic regions [45]. While acetylcholine receptor transcripts were largely confined to the NMJ in control muscle, denervation led to strong induction of multiple receptor subunits, including *Chrna1, Chrnb1*, and *Chrng*, throughout the myofiber (Fig. 5D). Notably, extrajunctional expression of *Chrnb1* and *Chrng* exceeded their baseline levels at the NMJ.

Importantly, not all receptor subunits exhibited the same spatial regulation. *Chrne* expression remained strictly confined to postsynaptic NMJ nuclei even after denervation (Fig. 5D), consistent with prior evidence that *Chrne* expression is developmentally regulated and controlled at individual endplates [46]. Spatial visualization further illustrated this dichotomy, showing persistent localization of postsynaptic markers such as *Prkar1a, Etv5*, and *Ufsp1* (red colors) at NMJs, alongside widespread induction of *Chrna1* and *Chrng* (yellow colors) across denervated muscle tissue (Fig. 5E). Together, these results demonstrate that denervation preserves postsynaptic NMJ identity while selectively derepressing extrajunctional acetylcholine receptor expression.

### Denervation Elicits Robust Extracellular Matrix Remodeling in the Perisynaptic NMJ Microenvironment

We next examined denervation-induced transcriptional changes in non-myofiber compartments associated with the NMJ microenvironment. In contrast to normal muscle (Fig. 3), transcriptomic signatures corresponding to myelinating Schwann cells, marked by *Mpz* and *Pmp22*, were not readily detected in denervated tissues. Nevertheless, perisynaptic regions defined by non-myelinating or terminal Schwann cell-associated markers remained identifiable in close proximity to postsynaptic NMJs (Fig. 5E).

Following denervation, perisynaptic marker expression became more spatially extensive compared to control muscle (Fig. 5E), suggesting expansion or activation of this compartment. Using marker-based segmentation analogous to that applied in normal muscle (Fig. 3), we isolated postsynaptic and perisynaptic NMJ regions and performed differential expression analysis between control and denervated conditions. This analysis revealed robust and selective upregulation of genes involved in extracellular matrix remodeling, tissue repair, and injury responses within the perisynaptic compartment (Fig. 5F-5I).

Spatial mapping confirmed that these extracellular matrix-associated transcripts were enriched in regions surrounding NMJs in denervated muscle (Fig. 5J). Given that many of these genes are expressed by fibroblasts and macrophages, it is possible that additional cell populations recruited to the NMJ following denervation contribute to the observed remodeling response. It is also possible that terminal Schwann cells underwent activation, proliferation, and extracellular matrix remodeling in response to neuronal injury [47–50]. Collectively, these results identify the perisynaptic NMJ microenvironment as a dynamic and transcriptionally active niche that undergoes pronounced extracellular matrix remodeling in response to denervation.

### Non-Myocyte Responses to Denervation

Finally, we examined transcriptional responses of macrophages and fibroblasts, two non-myocyte populations closely associated with myofiber integrity and tissue remodeling. In the hexagonal grid analysis (Fig. 1I; Fig. S1C-S1F), three subclusters were identified for each population, designated MACRO1-3 and FIBRO1-3 (Fig. S9A-S9I and Fig. S9J-S9S, respectively).

Across experimental conditions, macrophage populations showed a general trend toward increased abundance following denervation albeit with different kinetics (Fig. S9D-S9F), despite substantial variability in overall cell-type representation across samples (Fig. S1D-S1F). Concordantly, expression of macrophage-associated marker genes increased after denervation, with the highest levels observed at 7 days (Fig. S9G-S9I). These trends were evident across all three macrophage subclusters (MACRO1-3), indicating a broadly coordinated macrophage response rather than expansion of a single macrophage subtype.

In contrast, fibroblast populations did not exhibit a clear or consistent change in overall abundance following denervation (Fig. S9M-S9O). However, transcriptional profiling revealed pronounced condition-dependent remodeling within the FIBRO1 subcluster (Fig. S9J, S9M, S9P, S9S). Although the relative contribution of FIBRO1 cells remained largely stable at both 3 and 7 days after denervation (Fig. S9M), their gene expression profiles shifted substantially in a time-dependent manner.

At 3 days after denervation, FIBRO1 cells exhibited increased expression of genes associated with early injury and inflammatory responses, including the collagen *Col3a1*, the chemokine *Cxcl14*, and the lysosomal protease *Ctsl* (Fig. S9P, S9S). By 7 days, this transcriptional program transitioned toward enrichment of extracellular matrix-associated genes, including *Fn1, Mgp*, and *Dcn*, consistent with matrix remodeling processes (Fig. S9P, S9S). These expression patterns were reproducibly observed across individual samples, despite moderate inter-sample variability (Fig. S9S).

Together, these data indicate that non-myocyte populations respond to denervation through distinct and temporally structured transcriptional programs. Macrophages exhibit a progressive increase in abundance and activation, whereas fibroblasts, particularly the FIBRO1 subpopulation, undergo dynamic transcriptional remodeling without marked changes in population size. These findings highlight the importance of condition-dependent, cell type-specific gene regulation in shaping the non-myocyte contribution to denervation-induced muscle remodeling.

## Discussion

Skeletal muscle function is central to movement, metabolic homeostasis, and overall organismal health. A defining feature of this tissue is its pronounced cellular and subcellular heterogeneity, coupled with a strong dependence on neural input for maintenance of structure and function. Understanding how this complex architecture is organized in intact tissue and how it is remodeled following denervation is therefore essential for elucidating mechanisms underlying neuromuscular disease, aging-associated muscle decline, and recovery from nerve injury. Recent advances in transcriptomic technologies have created new opportunities to address these questions [51], yet each approach has inherent limitations that have constrained their application to skeletal muscle. In this study, we employed Seq-Scope, an ultra-high-resolution spatial transcriptomic platform, to generate a comprehensive atlas of normal and denervated mouse soleus muscle with single-cell and subcellular resolution.

To our knowledge, this work represents the first transcriptome-wide spatial atlas of skeletal muscle that combines microscopic spatial resolution with unbiased gene coverage in longitudinal sections. Previous transcriptomic studies of skeletal muscle have relied primarily on scRNA-seq or snRNA-seq, which necessarily disrupt tissue architecture and eliminate spatial context, or on spatial platforms such as 10X Visium, whose limited resolution precludes reliable discrimination of individual myofibers or subcellular domains [15–22]. As a result, prior spatial transcriptomic studies of skeletal muscle have been restricted to transverse sections and were unable to assess transcriptomic variation along the longitudinal axis of myofibers. More recently, imaging-based approaches such as 10X Xenium have achieved cellular-scale resolution in skeletal muscle [52]; however, these methods rely on targeted gene panels and therefore capture only a small fraction (<400 genes) of the transcriptome. By contrast, Seq-Scope provides submicrometer spatial resolution together with whole-transcriptome coverage, enabling direct visualization of transcriptional organization within intact myofibers and across tissue microenvironments in longitudinal sections.

To extract biological insight from the resulting high-dimensional dataset, we employed two complementary analytical strategies: histology-guided myofiber segmentation and unbiased hexagonal grid-based spatial binning. These approaches produced convergent yet distinct results. Segmentation-based analysis enabled robust classification of major myofiber types and supported comprehensive fiber type-specific differential gene expression analyses. In contrast, grid-based spatial analysis localized transcriptomic phenotypes without reliance on predefined cellular boundaries, revealing hybrid myofiber states, subcellular transcriptional domains, and localized degenerative signatures that were not resolved by segmentation alone. Importantly, this approach also captured spatially organized non-myocyte populations, including macrophages, fibroblasts, smooth muscle cells, and adipocytes, as well as specialized structures such as neuromuscular junctions and partially degenerating myofiber regions. Together, these complementary strategies provide a holistic molecular view of skeletal muscle architecture across the longitudinal histological axis.

Our results further underscore the complexity of denervation-induced remodeling in skeletal muscle. At the level of myofibers, denervation elicited robust and fiber type-specific transcriptional responses, most prominently at early time points. These responses included induction of stress-associated pathways involving heat shock proteins, crystallins, metallothioneins, autophagy regulators, and genes related to oxygen metabolism. Notably, these transcriptional changes were not uniformly distributed along myofibers. Instead, high-resolution spatial analysis revealed localized domains of damage-response gene expression, frequently enriched near neuromuscular junctions and associated with histological features of partial degeneration. Such spatially restricted responses are difficult to detect using dissociation-based approaches or low-resolution spatial methods [15–22] and are not accessible through targeted imaging-based assays [52].

Non-myocyte populations also exhibited distinct and temporally structured responses to denervation. Macrophages showed a progressive increase in abundance and expression of macrophage-associated genes, consistent with their established roles in injury response and tissue remodeling. Fibroblasts, in contrast, did not display a consistent change in population size but instead underwent pronounced transcriptional remodeling within a defined subpopulation. Early denervation was associated with increased expression of collagens, chemokines, and proteases, whereas later stages were characterized by upregulation of extracellular matrix components, suggesting a transition from early injury-associated signaling to matrix remodeling. These findings indicate that denervation-induced changes in non-myocyte compartments are driven largely by dynamic gene regulation rather than simple shifts in cellular abundance.

At the neuromuscular junction, Seq-Scope resolved coordinated yet distinct transcriptional responses across postsynaptic, perisynaptic, and Schwann cell-associated domains. Denervation preserved the transcriptional identity of postsynaptic NMJ nuclei while inducing widespread extrajunctional expression of select acetylcholine receptor subunits across myofibers, consistent with established activity-dependent regulation of these genes [45, 46]. In parallel, perisynaptic regions exhibited expansion and robust induction of extracellular matrix and injury-response genes, in agreement with previous reports describing activation of terminal Schwann cells and associated remodeling processes following nerve injury [36, 47–50]. The ability to spatially disentangle these compartment-specific responses highlights the strength of high-resolution spatial transcriptomics for dissecting multicellular niches within intact tissue.

In conclusion, this study provides a detailed spatial and transcriptional framework for understanding skeletal muscle organization and its remodeling following denervation. By leveraging the resolution and transcriptome coverage of Seq-Scope, we uncover heterogeneity across fiber types, within individual myofibers, and among diverse non-myocyte populations, revealing spatially structured responses that are not accessible with previous transcriptomic approaches. While these findings are derived from a murine model and will require validation in human tissue and disease contexts, this atlas establishes a reference for future studies of neuromuscular pathology, aging, and regeneration. More broadly, it demonstrates the power of high-resolution spatial transcriptomics to advance understanding of complex, multinucleated tissues and their dynamic responses to physiological perturbation.

## Data Availability

All data from this study have been deposited in the Deep Blue Data repository (https://doi.org/10.7302/8w29-zc33) and will be made available to the community upon provisional acceptance of the paper.

## Methods

### Mice Muscle Injury Model

All animal procedures were subject to the regulations established by the University of Michigan Institutional Animal Care and Use Committee (IACUC). C57BL/6J mice littermates, approximately 6 months of age, underwent sciatic nerve transection (SNT) on their left hindlimb and were randomly assigned to a 3-day or 7-day recovery group. Mice were anesthetized using 5% isoflurane (MWI Animal Health) followed by continuous 2% isoflurane administration to ensure an unresponsive state to tactile stimuli. Preoperative analgesia was administered using 5 mg/kg carprofen (MWI Animal Health). The hindlimbs were shaved and cleaned with chlorhexidine and 70% alcohol. A small incision (<10 mm) was made 1 mm posterior and parallel to the femur to expose the sciatic nerve. The sciatic nerve was transected with a removal of a 5 mm segment, and the incision was closed with wound clips (Autoclip, BD Clay Adams). A sham procedure was replicated on the contralateral leg. Mice were placed on a heating pad after the procedure and monitored until full recovery from anesthesia. Mice recovered for either three or seven days before being sacrificed and tissue collection.

### Seq-Scope array construction (1st-Seq)

Seq-Scope array was constructed as described previously [24]. Seq-Scope array is a solid phase transcriptome capturing system, which was constructed on the HiSeq2000 (Illumina) platform with HDMI32-DraI, a single-stranded oligonucleotide library (Eurofins), and Read1-DraI sequencing primer. Through sequence-by-synthesis strategy, 100pM HDMI32-DraI oligonucleotides were placed and formed into clusters on the flow cell surface. Sequencer was manually set to perform 37bp single-end reading with a custom Read1-DraI primer. The run was completed right after 37 cycles, and the flow cell was retrieved. The output FASTQ file from the run will provide the sequence and the XY coordinates of the barcodes. The retrieved flow cell was further processed into ready-to-use Seq-Scope array. The flow cell was washed with nuclease-free water three times and incubated with DraI (R0129, NEB) and CIAP (M0525, NEB) enzyme mixture in 37 °C overnight to expose oligo-dT end. The flow cell was then treated with exonuclease I (M2903, NEB) cocktail in 37 °C for 45 min to remove non-specific single-stranded DNA. In the last step, the flow cell underwent a series of washing steps, including three times with nuclease-free water, three times with 0.1N NaOH for 5 min each, and three times with 0.1M Tris pH7.5. The cover of the flow cell was then disassembled using Tungsten Carbide Tip Scriber (IMT-8806, IMT) to expose the surface of the array for tissue attachment.

### Tissue Preparation, Sectioning, and processing

Mice subjected to SNT were anesthetized through intraperitoneal injection of 0.5 mg/g avertin (2,2,2,-Tribromoethanol 97%, Sigma #T48402, and 2-methyl-2-butanol 99%, Sigma #240486-100ml). Soleus muscles were freshly dissected and embedded into O.C.T compound (23-730-571, Fisher Scientific). The OCT blocks were snap-frozen by submerging into liquid nitrogen pre-chilled 2-Methylbutane (MX0760, Sigma-Aldrich). The sectioning of the OCT-mounted frozen tissue was performed in a cryostat (Leica CM3050S, -15 °C) at a 5° cutting angle with each section 10 μm thick. Sections were attached to the array and re-warmed in room temperature. Attached sections were then fixed in 4% formaldehyde (15170, Electron Microscopy Sciences) in room temperature for 10 min, followed by hematoxylin and eosin (H&E) staining that was described previously [24]. H&E images were captured under the Keyence digital darkroom system.

### Seq-Scope RNA Library Generation and Sequencing (2nd-Seq)

Seq-Scope library construction was constructed as described previously [24]. In brief, tissue sections were permeabilized with 0.2 U/μL collagenase I (17018-029, Thermo Fisher) at 37 °C for 20 min, followed with 1mg/mL pepsin (P7000, Sigma) in 0.1M HCl at 37 °C for 10 min. Permeabilized tissue sections were then washed with 1X Maxima RT buffer (EP0751, Thermofisher). Reverse transcription was conducted on the array through the incubation of RT mixture (1X RT buffer (EP0751, Thermofisher), 4% Ficoll PM-400 (F4375-10G, Sigma), 1mM dNTPs (N0477L, NEB), RNase inhibitor (30281, Lucigen), Maxima H-RTase (EP0751, Thermofisher)) in a humidified chamber at 42 °C overnight.

Tissue sections were treated with exonuclease I (#M2903, NEB) cocktail the next day in 37 °C for 45 min to remove unused single-stranded DNA probes. Sections were then incubated in tissue digestion cocktail (100 mM Tris pH 8.0, 100 mM NaCl, 2% SDS, 5 mM EDTA, 16 U/mL Proteinase K (P8107S, NEB)) at 37 °C for 40 min. After the tissues were digested, the array was washed three times with nuclease-free water, three times with 0.1N NaOH for 5 min each, and three times with 0.1M Tris pH7.5.

Secondary stand synthesis was conducted following the washing step. The array was loaded with secondary strand synthesis mixture (1X NEBuffer-2 (NEB), 10 uM TruSeq Read2-conjugated Random Primer (IDT), 1mM dNTPs (N0477, NEB), Klenow Fragment (M0212, NEB), and nuclease-free water) and incubated in a humidified chamber at 37 °C for 2 hr. Solution was removed upon the completion of the reaction, and the array was washed with nuclease-free water three times. The array was then treated with 0.1N NaOH for 5 min to elute the barcoded cDNA library. The pH of the elution was neutralized by 3 M potassium acetate, pH5.5. Eluted library was then further purified using AMPure XP beads (1.8X bead/sample ratio, A63881, Beckman Coulter) according to the manufacturer’s instruction.

Library amplification was conducted through two rounds of PCR. First round PCR was performed using Kapa HiFi Hotstart Readymix (KK2602, KAPA Biosystems), 2μM forward and reverse primers as described in our previous paper [24], and the total eluted library. PCR product was purified using AMPure XP beads (A63881, Beckman Coulter) with 1.2X beads/sample ratio. Second round PCR was performed using Kapa HiFi Hotstart Readymix (KK2602, KAPA Biosystems), indexing primers described in [24], and 2nM of the first round PCR product. PCR product was purified again using AMPure XP beads (A63881, Beckman Coulter) with 0.6X beads/sample ratio. The size of the library was selected between 400-850bp through agarose gel electrophoresis. Selected library was purified using Zymoclean Gel DNA Recovery Kit (D4001, Zymo Research). The library was then sequenced with a paired-end (100-150bp) setup.

### Generation of Spatial Digital Gene Expression (sDGE) Matrix

Data are processed as described previously [24, 28], combing both 1st-Seq and 2nd-Seq FASTQ results. In the processing, 2nd-Seq FASTQ results were first filtered for their presence in 1st-Seq data table, so that the alignment and data processing could be more efficient. Then alignment was performed in STAR to produce spatial digital gene expression matrix (sDGE) matching individual gene expression with spatial coordinates identified through HDMI matching between 1st-Seq and 2nd-Seq.

### Alignment between histology and spatial dataset

From the dataset, spatial transcriptome density is visualized through hillshade function. The hillshade image was semi-automatically aligned with H&E image using QGIS (version 3.22.9) georeferencer function, through manually marking salient spatial features common in both images.

### Myofiber Segmentation and Hexagonal Gridding

Myofiber segmentation was performed on QGIS by manually drawing polygons on H&E images, according to myofiber boundary lines. Transcripts within each fiber were aggregated into a single datapoint, which was treated as a single cell located at a point on surface of the polygon. For hexagonal gridding, spatial data are aggregated toward a non-overlapping array of hexagons with 14µm sides, which was treated as a single cell located at the center of each hexagon.

### Data Analysis and Visualization

The aggregated DGE matrix was analyzed in the Seurat package [53]. Feature number threshold was applied to remove the grids (as indicated in the figure and the text) that corresponded to the area that was not overlaid by the tissue. Data were normalized using SCTransform function. After discovering principal components through RunPCA and generating UMAP manifold using RunUMAP, clustering was performed using the shared nearest neighbor modularity optimization implemented in Seurat’s FindNeighbors and FindClusters function, and then annotated to specific myofiber and cell types based on marker gene expression. For clustering segmented myofiber transcriptome from control/denervated dataset, clustering was sequentially performed, first to differentiate experimental conditions, and then to identify myofiber subtypes in each of experimental condition-dependent clusters. Clusters were visualized in histological space through plotting hexagonal centers using ggplot2 package (hexagonal dataset) or through plotting the whole polygonal area using geopandas and matplotlib package (segmentation dataset). FindMarkers and FindAllMarkers functions were used to identify Top markers from each cluster, and volcano plot was generated using the output of these functions using ggplot2.

Level of gene expression was visualized though Seurat functions such as VlnPlot, DotPlot and FeaturePlot, and spatial gene expression was visualized through a custom visualization software which visualizes discovered genes as colored dots in the coordinate space. Area-proportional Venn diagrams were made using DeepVenn [54].

Overlapping hexagonal grid were generated to allow for multiscale sliding windows analysis [24, 28]. To make a projection, annotations provided from non-overlapping hexagonal analysis were fine tuned using FindTransferAnchors and TransferData in Seurat. Separately, factors learned from clusters of non-overlapping hexagonal analysis were projected into raw pixel-level dataset through FICTURE as described in our recent work [55].

---

**Fig. S1.**
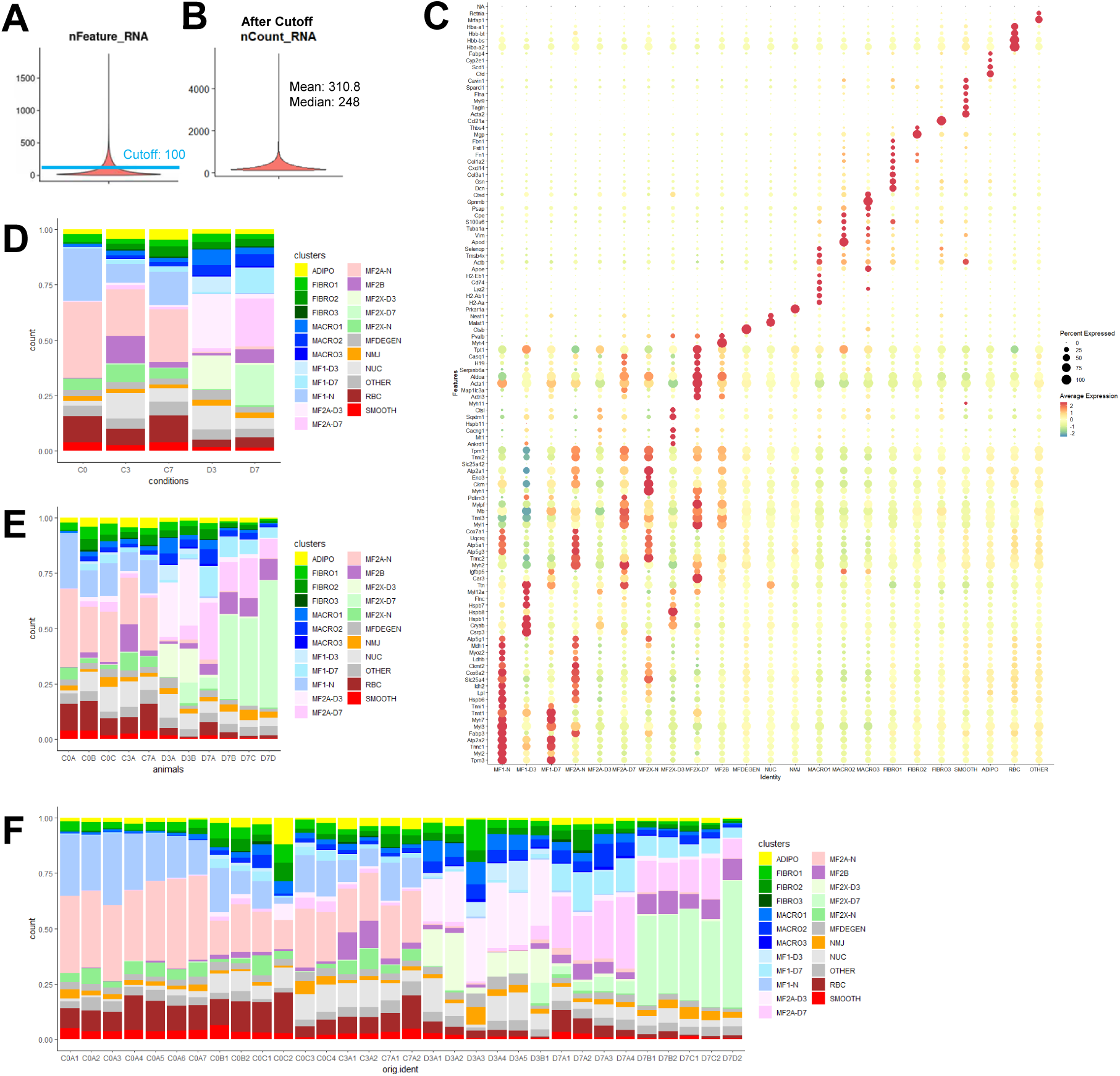
Transcriptome-based identification of cell types from Seq-Scope data. (A, B) Myofiber spatial transcriptomes were segmented into hexagonal grids with 14 µm side length and 24 µm flat-to-flat height. (A) Violin plot showing the distribution of gene features per hexagon (nFeature_RNA). A cutoff of 100 gene features (blue line) was applied to obtain high-quality transcriptomes. (B) Violin plot showing the distribution of unique transcript counts per hexagon (nCount_RNA) after applying the 100 gene feature cutoff. (C) Expression of cluster-specific marker genes used for cell type mapping shown in Fig. 1F and Fig. 1I. (D-F) Distribution of identified clusters across experimental conditions (D), animal identity (E), and section identity (F). Individual section identities in (F) are encoded by four letters, with the first two letters indicating experimental condition as in Fig. 1A and the third and fourth letters denoting animal and section identity, respectively.

**Fig S2.**
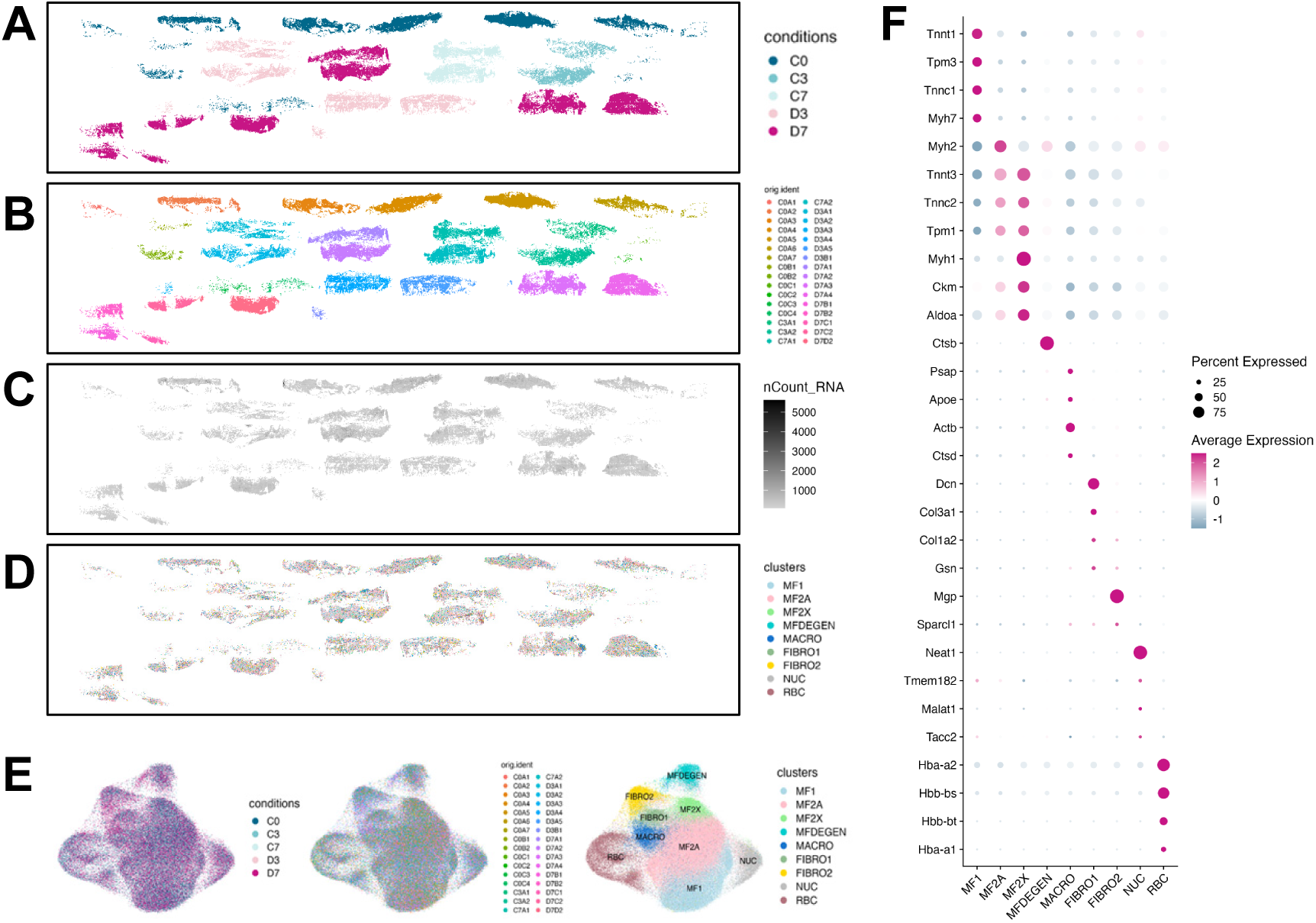
Data integration identifies canonical myofiber types and major muscle cell populations. (A-F) Spatial distribution of experimental conditions (A), individual section identities (B), unique transcript density (C), and transcriptome clustering results (D) following integration (E). In (E), UMAP embeddings of the integrated dataset are colored by experimental condition (left), individual section identity (middle), and cluster assignment (right), demonstrating effective batch correction. Expression of cell type markers across integrated clusters is shown in a dot plot (F). Individual section identities in (B, E) are encoded by four letters, with the first two letters denoting experimental condition as in Fig. 1A and the third and fourth letters indicating animal and section identity, respectively.

**Fig. S3.**
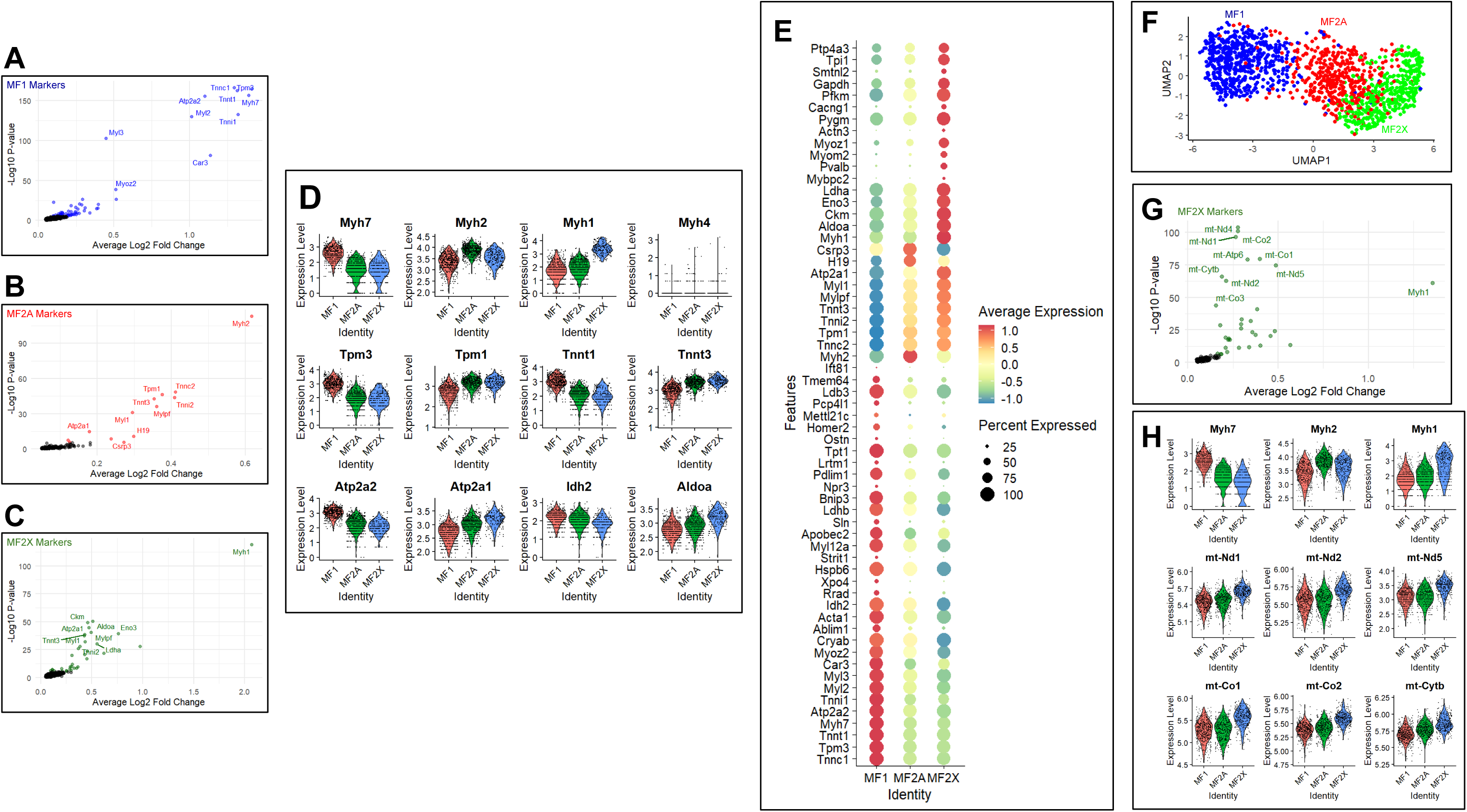
Fiber type-specific gene expression in single segmented myofibers. (A-C) Volcano plots showing genes enriched in individual myofiber clusters, MF1 (A), MF2A (B), and MF2X (C). The top 20 marker genes are highlighted using the same color scheme as in Fig. 2E. (D) Violin plots illustrating expression of selected genes across myofiber populations, including myosin heavy chain genes (top), tropomyosins and troponins (middle), and endoplasmic reticulum calcium pumps and metabolic enzymes (bottom). (E) Dot plot showing expression of additional marker genes enriched in the myofiber clusters identified in Fig. 2E. (F) UMAP representation of the myofiber transcriptome derived from a dataset that includes mitochondrial and hypothetical genes. The clustering pattern is largely consistent with the analysis excluding these genes shown in Fig. 2E. (G) Volcano plot identifying marker genes enriched in MF2X fibers from (F), highlighting elevated expression of mitochondrial genes. (H) Violin plots showing expression patterns of myosin heavy chain genes and selected mitochondrially encoded genes across the myofiber populations identified in (F).

**Fig. S4.**
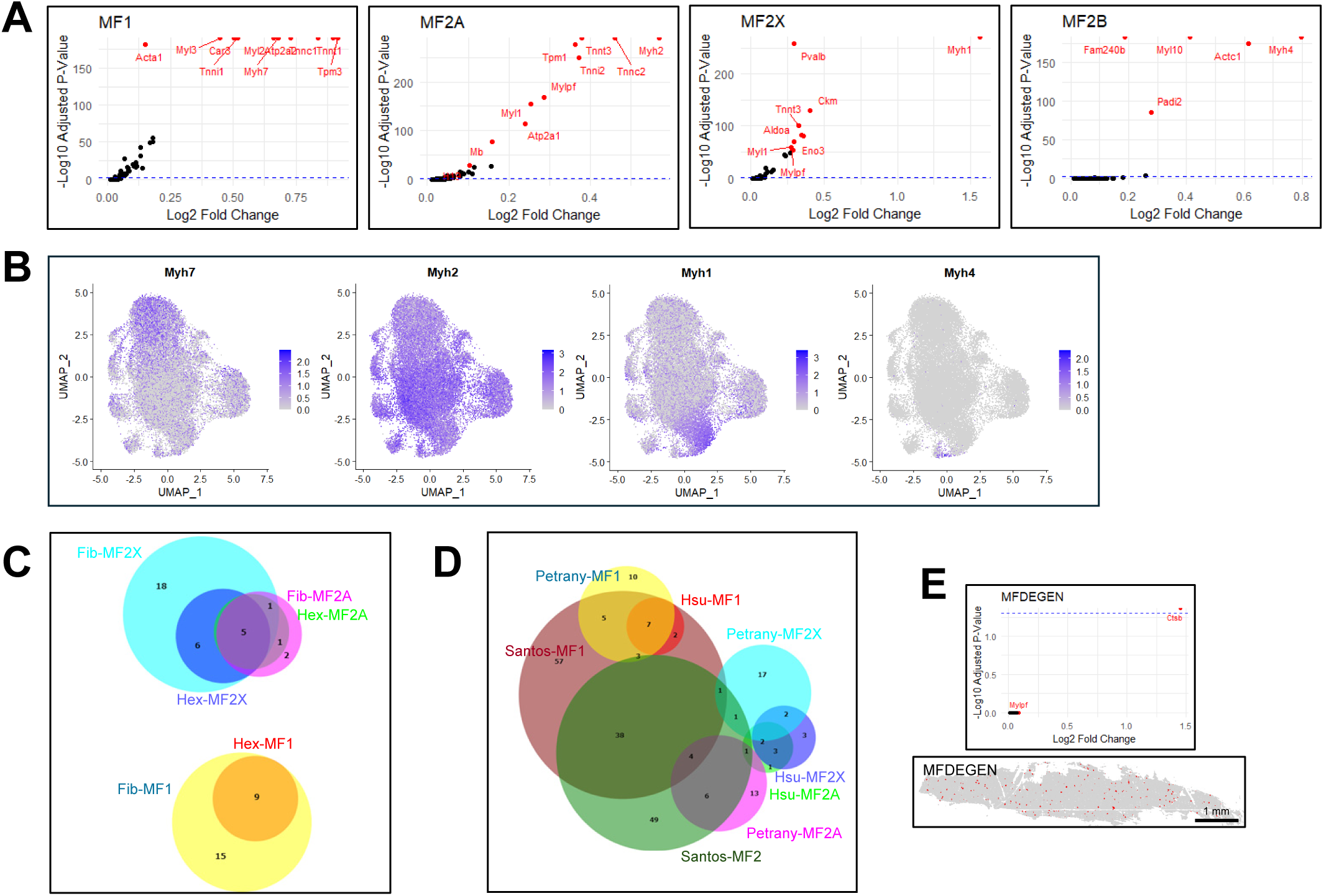
Characterization of myofiber type transcriptomes using hexagonal grid analysis. (A) Volcano plots showing genes upregulated in the indicated myofiber types. (B) Expression of fiber type-specific myosin heavy chain isoforms projected onto the UMAP manifold. (C) Comparison of marker gene sets for each myofiber transcriptome between the hexagonal grid-based analysis (Hex) shown in Fig. 2G-H and the segmentation-based analysis (Fib) shown in Fig. 2E, using Venn diagrams. (D) Comparison of marker gene sets identified in the hexagonal grid-based myofiber transcriptomes with those reported in two previous single-nucleus RNA sequencing studies [7, 8], using a Venn diagram generated with DeepVenn [54]. (E) Volcano plot identifying marker genes enriched in MFDEGEN cluster (top) and their spatial distribution in histological space (bottom).

**Fig. S5.**
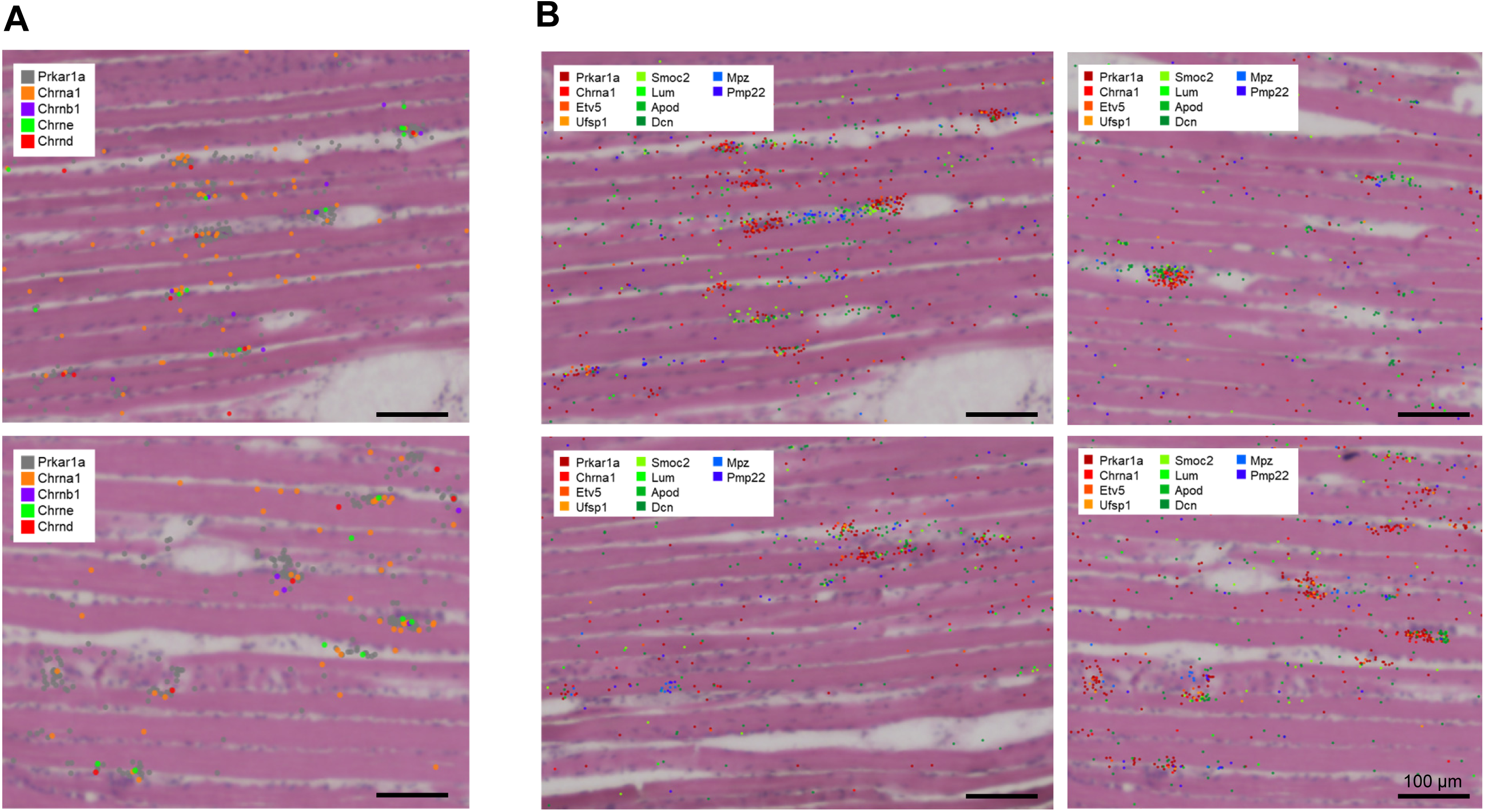
Expression of distinct marker genes in the neuromuscular junction region. (A) Spatial expression of acetylcholine receptor isoforms around postsynaptic nuclei identified by *Prkar1a* expression. (B) Spatial expression of cell type-specific marker genes identified in the analysis shown in Fig. 3 across distinct NMJ-associated regions.

**Fig. S6.**
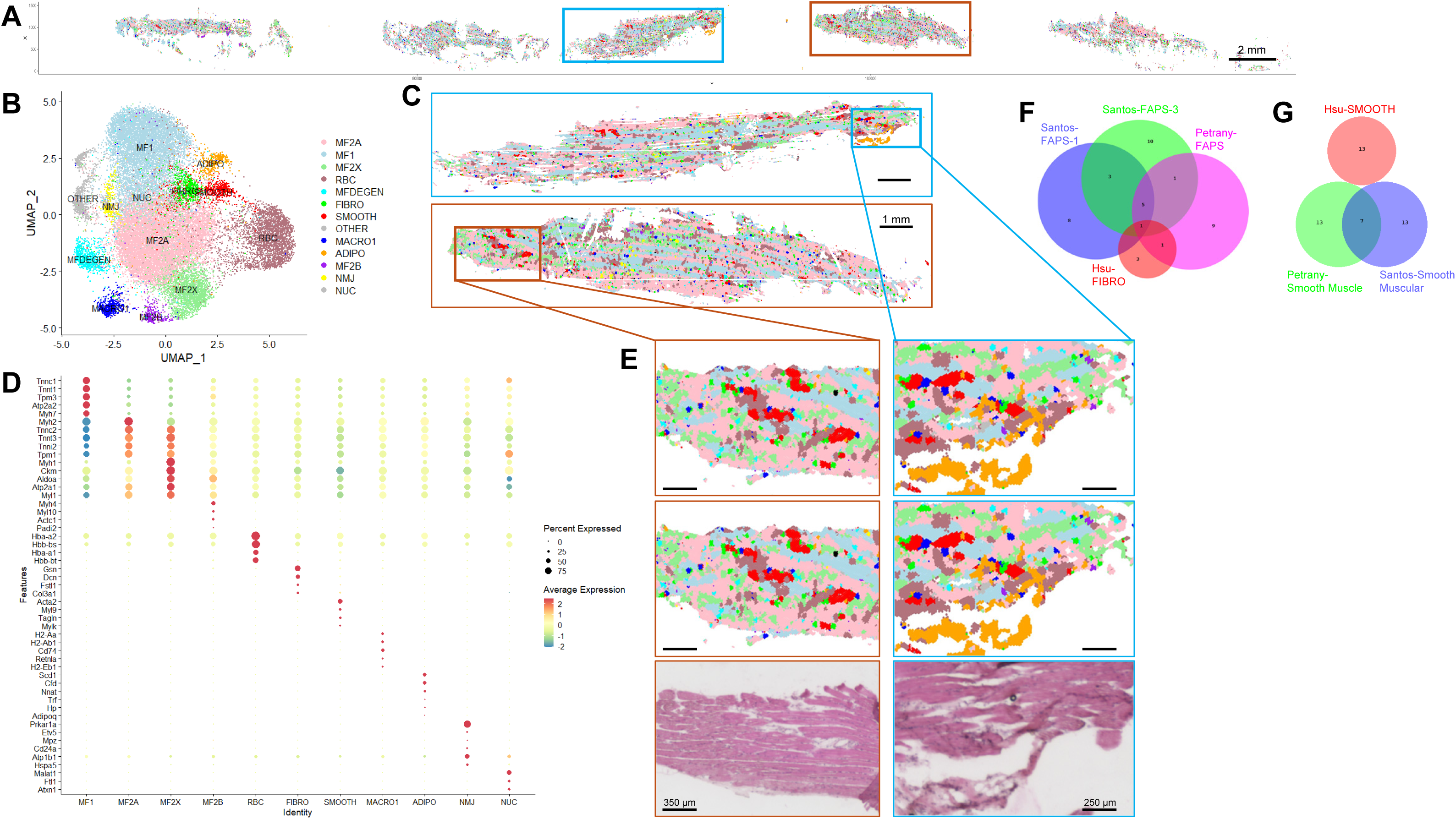
Non-myocyte transcriptomic heterogeneity revealed in the Seq-Scope dataset. (A, B) Spatial map (A) and UMAP manifold (B) from Fig. 2G and Fig. 2H, recolored to emphasize non-myocyte transcriptomes. (C) Magnified spatial views from the boxed regions in (A) shown with corresponding cluster colors. (D) Dot plots showing expression of top cluster-specific marker genes across non-myocyte clusters. (E) Further magnification of boxed regions in (C) as indicated by guide lines. Left panels highlight arteries, while right panels highlight adipose tissue. (F, G) Comparison of marker gene sets for fibroblasts or fibro-adipogenic progenitors (FAPs) (F) and smooth muscle cells (G) between the current hexagonal analysis and two previous single-nucleus RNA sequencing studies [7, 8]. The top 20 significant genes were included in the analysis.

**Fig. S7.**
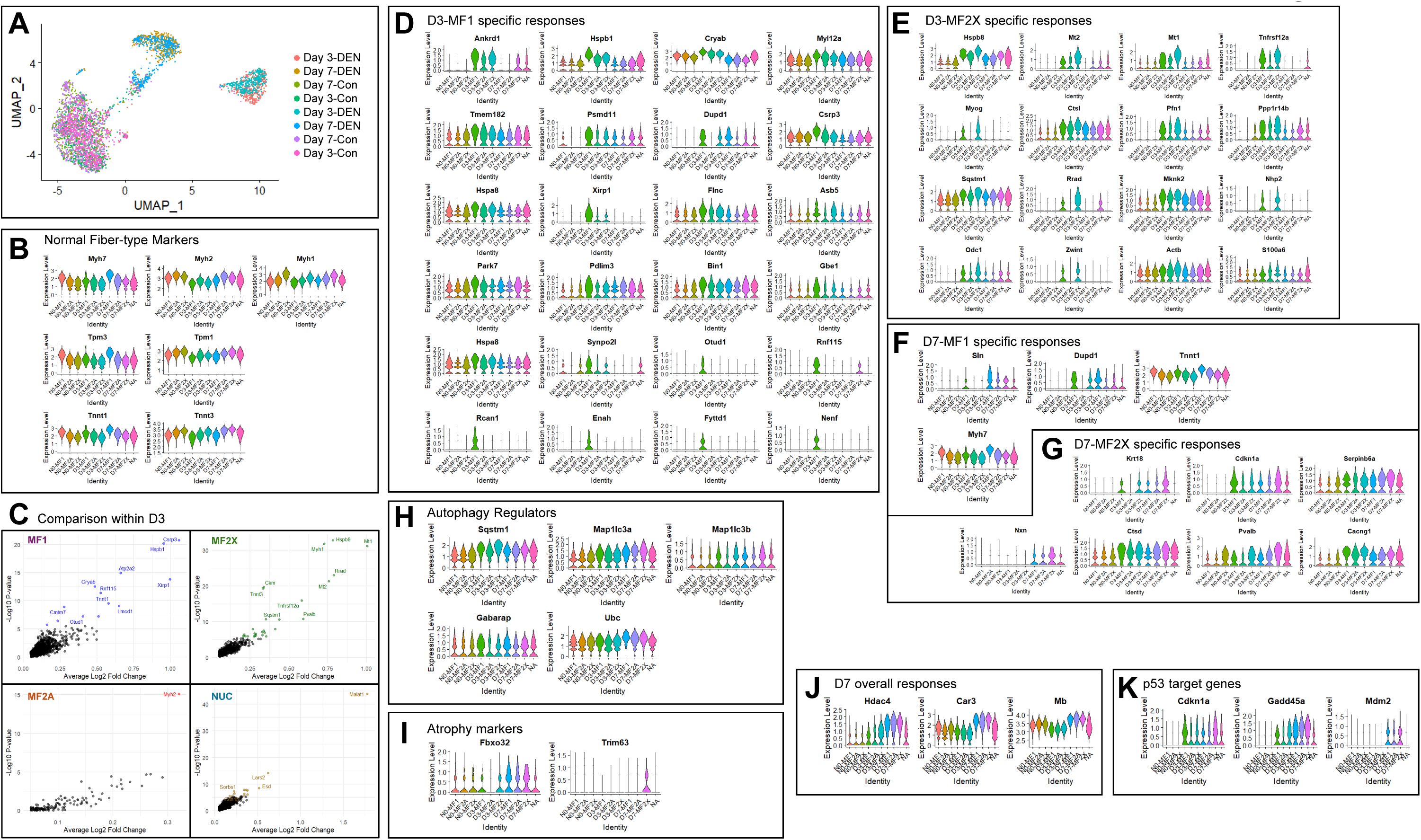
Fiber type-specific denervation responses. (A) UMAP manifold of the myofiber transcriptome colored by experimental condition. Control samples integrate well, whereas day 3 and day 7 denervated samples form distinct clusters corresponding to their respective time points. (B) Expression of myosin heavy chain isoforms, tropomyosins, and troponins across myofiber types in the denervated segmented myofiber dataset. (C) Identification of myofiber type-specific denervation response genes. Genes highly expressed in each myofiber type within the day 3 denervated dataset are highlighted. (D-K) Gene induction across myofiber types in the denervated dataset, grouped as day 3 myofiber type I-specific responses (D), day 3 myofiber type IIx-specific responses (E), day 7 myofiber type I-specific responses (F), day 7 myofiber type IIx-specific responses (G), autophagy-related responses (H), atrophy-related responses (I), day 7 global responses (J), and DNA damage responses (K).

**Fig. S8.**
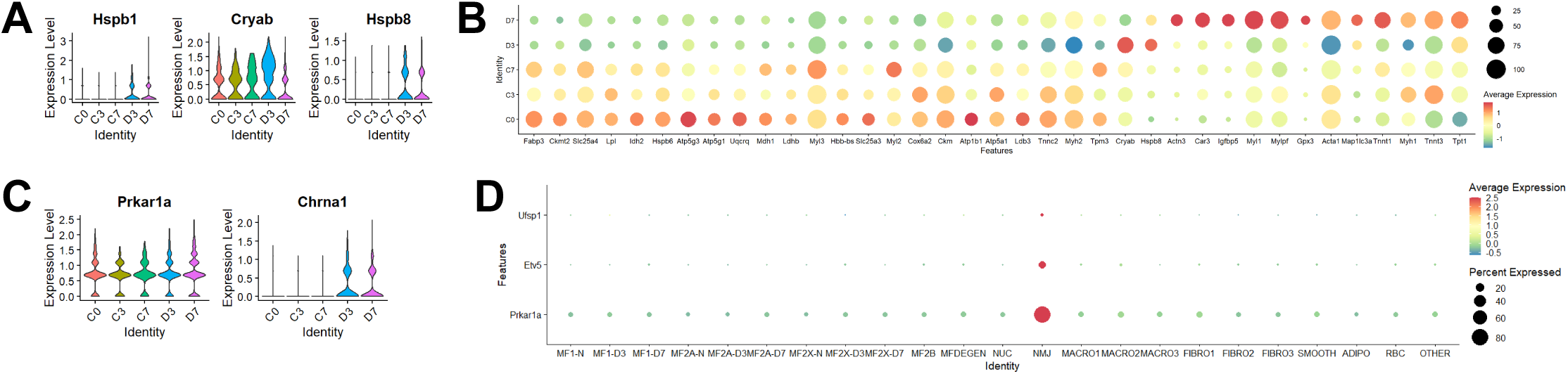
Neuromuscular junction responses to denervation. (A-C) Expression of the indicated genes across different experimental conditions, visualized by ViolinPlot (A, C) and DotPlot (B). (D) Differential expression of postsynaptic nuclei marker genes across different clusters, visualized by DotPlot analysis.

**Fig. S9.**
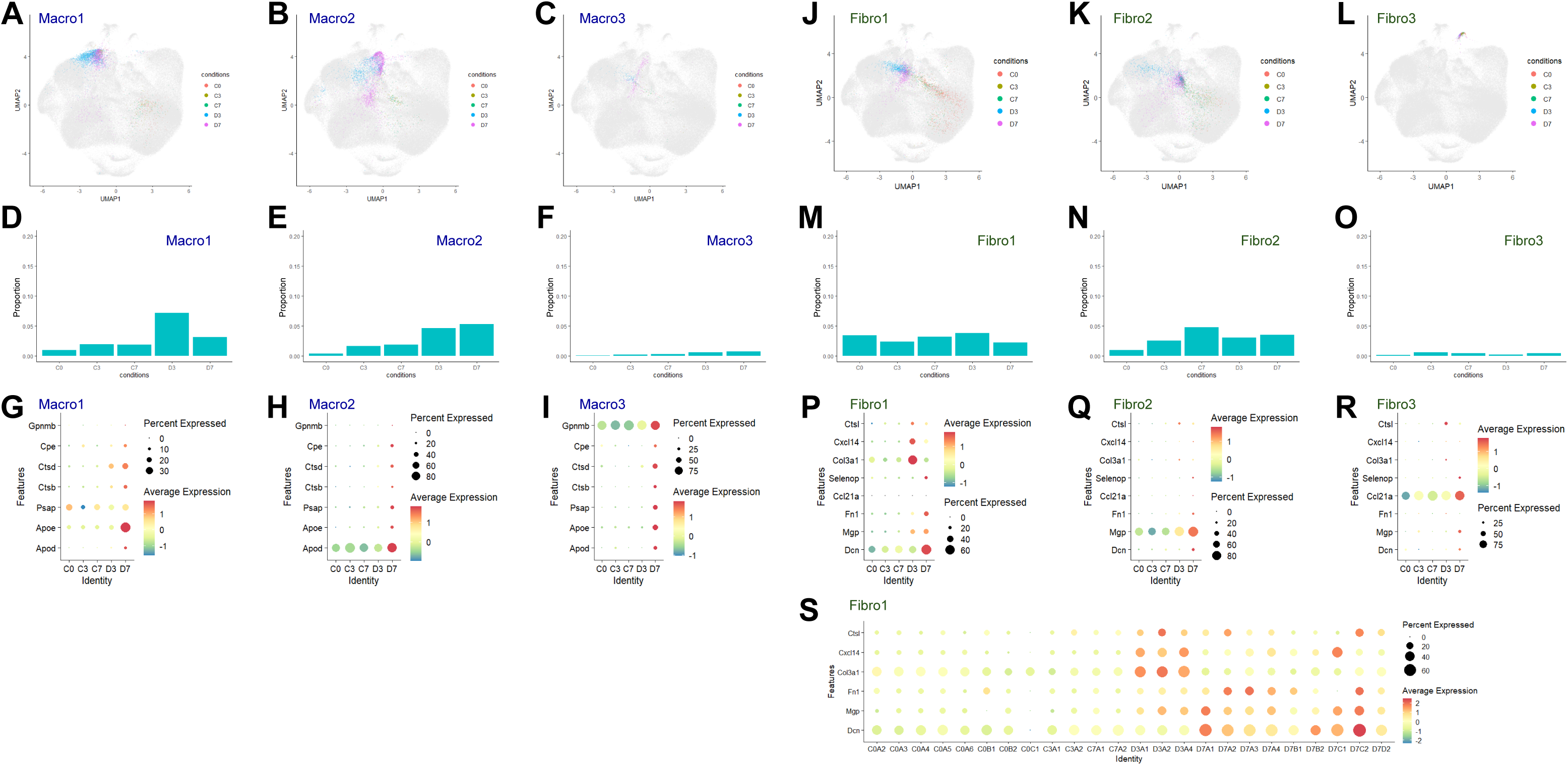
Denervation responses in non-myocytes. (A-C) UMAP manifold from Fig. 1I highlighting the indicated macrophage clusters, colored by experimental condition. All other clusters are shown in gray in the background. (D-F) Population shares of the indicated macrophage clusters across experimental conditions. (G-I) Differential expression of cluster-specific marker genes in macrophage subclusters across experimental conditions. (J-L) UMAP manifold from Fig. 1I highlighting the indicated fibroblast clusters, colored by experimental condition. All other clusters are shown in gray in the background. (M-O) Population shares of the indicated fibroblast clusters across experimental conditions. (P-R) Differential expression of cluster-specific marker genes in fibroblast subclusters across experimental conditions. (S) Expression of the indicated cluster-specific marker genes in fibroblast subclusters across individual samples.

## References

1. Mukund, K. and S. Subramaniam, Skeletal muscle: A review of molecular structure and function, in health and disease. Wiley Interdiscip Rev Syst Biol Med, 2020. 12(1): p. e1462.

2. Ritso, M., L.W. Tung, and F.M.V. Rossi, Emerging skeletal muscle stromal cell diversity: Functional divergence in fibro/adipogenic progenitor and mural cell populations. Exp Cell Res, 2022. 410(1): p. 112947.

3. Schiaffino, S. and C. Reggiani, Fiber types in mammalian skeletal muscles. Physiol Rev, 2011. 91(4): p. 1447–531.

4. Williams, K., K. Yokomori, and A. Mortazavi, Heterogeneous Skeletal Muscle Cell and Nucleus Populations Identified by Single-Cell and Single-Nucleus Resolution Transcriptome Assays. Front Genet, 2022. 13: p. 835099.

5. Ehmsen, J.T. and A. Hoke, Cellular and molecular features of neurogenic skeletal muscle atrophy. Exp Neurol, 2020. 331: p. 113379.

6. Soendenbroe, C., J.L. Andersen, and A.L. Mackey, Muscle-nerve communication and the molecular assessment of human skeletal muscle denervation with aging. Am J Physiol Cell Physiol, 2021. 321(2): p. C317–C329.

7. Petrany, M.J., et al., Single-nucleus RNA-seq identifies transcriptional heterogeneity in multinucleated skeletal myofibers. Nat Commun, 2020. 11(1): p. 6374.

8. Dos Santos, M., et al., Single-nucleus RNA-seq and FISH identify coordinated transcriptional activity in mammalian myofibers. Nat Commun, 2020. 11(1): p. 5102.

9. Kim, M., et al., Single-nucleus transcriptomics reveals functional compartmentalization in syncytial skeletal muscle cells. Nat Commun, 2020. 11(1): p. 6375.

10. Orchard, P., et al., Human and rat skeletal muscle single-nuclei multi-omic integrative analyses nominate causal cell types, regulatory elements, and SNPs for complex traits. Genome Res, 2021. 31(12): p. 2258–2275.

11. Lin, H., et al., Decoding the transcriptome of denervated muscle at single-nucleus resolution. J Cachexia Sarcopenia Muscle, 2022. 13(4): p. 2102–2117.

12. Tian, L., F. Chen, and E.Z. Macosko, The expanding vistas of spatial transcriptomics. Nat Biotechnol, 2023. 41(6): p. 773–782.

13. Bressan, D., G. Battistoni, and G.J. Hannon, The dawn of spatial omics. Science, 2023. 381(6657): p. eabq4964.

14. Kang, H.M. and J.H. Lee, Spatial Single-Cell Technologies for Exploring Gastrointestinal Tissue Transcriptome. Compr Physiol, 2023. 13(3): p. 4709–4718.

15. D’Ercole, C., et al., Spatially resolved transcriptomics reveals innervation-responsive functional clusters in skeletal muscle. Cell Rep, 2022. 41(12): p. 111861.

16. Heezen, L.G.M., et al., Spatial transcriptomics reveal markers of histopathological changes in Duchenne muscular dystrophy mouse models. Nat Commun, 2023. 14(1): p. 4909.

17. Larouche, J.A., et al., Spatiotemporal mapping of immune and stem cell dysregulation after volumetric muscle loss. JCI Insight, 2023. 8(7).

18. Ruoss, S., et al., Spatial transcriptomics tools allow for regional exploration of heterogeneous muscle pathology in the pre-clinical rabbit model of rotator cuff tear. J Orthop Surg Res, 2022. 17(1): p. 440.

19. Stec, M.J., et al., A cellular and molecular spatial atlas of dystrophic muscle. Proc Natl Acad Sci U S A, 2023. 120(29): p. e2221249120.

20. Patsalos, A., et al., Spatiotemporal transcriptomic mapping of regenerative inflammation in skeletal muscle reveals a dynamic multilayered tissue architecture. J Clin Invest, 2024. 134(20).

21. Coulis, G., et al., Single-cell and spatial transcriptomics identify a macrophage population associated with skeletal muscle fibrosis. Sci Adv, 2023. 9(27): p. eadd9984.

22. Lequain, H., et al., Spatial Transcriptomics Reveals Signatures of Histopathological Changes in Muscular Sarcoidosis. Cells, 2023. 12(23).

23. Gurkar, A.U., et al., Spatial mapping of cellular senescence: emerging challenges and opportunities. Nat Aging, 2023. 3(7): p. 776–790.

24. Cho, C.-S., et al., Microscopic examination of spatial transcriptome using Seq-Scope. Cell, 2021. 184(13): p. 3559–3572. e22.

25. Pette, D. and C. Spamer, Metabolic properties of muscle fibers. Fed Proc, 1986. 45(13): p. 2910–4.

26. Do, T.H., et al., TREM2 macrophages induced by human lipids drive inflammation in acne lesions. Sci Immunol, 2022. 7(73): p. eabo2787.

27. Si, Y., et al., FICTURE: scalable segmentation-free analysis of submicron-resolution spatial transcriptomics. Nat Methods, 2024. 21(10): p. 1843–1854.

28. Xi, J., et al., STtools: a comprehensive software pipeline for ultra-high-resolution spatial transcriptomics data. Bioinformatics Advances, 2022. 2(1).

29. Gokce, E. and N. Gun, The Relationship Between Exercise, Cathepsin B, and Cognitive Functions: Systematic Review. Percept Mot Skills, 2023. 130(4): p. 1366–1385.

30. Kar, N.C. and C.M. Pearson, Early elevation of cathepsin B1 in human muscle disease. Biochem Med, 1977. 18(1): p. 126–9.

31. West, J.A., et al., The long noncoding RNAs NEAT1 and MALAT1 bind active chromatin sites. Mol Cell, 2014. 55(5): p. 791–802.

32. Wolbert, J., et al., Redefining the heterogeneity of peripheral nerve cells in health and autoimmunity. Proc Natl Acad Sci U S A, 2020. 117(17): p. 9466–9476.

33. Pantera, H., et al., Regulation of the neuropathy-associated Pmp22 gene by a distal super-enhancer. Hum Mol Genet, 2018. 27(16): p. 2830–2839.

34. Berger, P., et al., Expression analysis of the N-Myc downstream-regulated gene 1 indicates that myelinating Schwann cells are the primary disease target in hereditary motor and sensory neuropathy-Lom. Neurobiol Dis, 2004. 17(2): p. 290–9.

35. Castro, R., et al., Specific labeling of synaptic schwann cells reveals unique cellular and molecular features. Elife, 2020. 9.

36. Guzman, S.D., et al., Decoding muscle-resident Schwann cell dynamics during neuromuscular junction remodeling. JCI Insight, 2025.

37. Seaberg, B.L., S. Purao, and M. Rimer, Validation of terminal Schwann cell gene marker expression by fluorescent in situ hybridization using RNAscope. Neurosci Lett, 2022. 771: p. 136468.

38. Zhang, S., et al., Fibroblastic SMOC2 Suppresses Mechanical Nociception by Inhibiting Coupled Activation of Primary Sensory Neurons. J Neurosci, 2022. 42(20): p. 4069–4086.

39. Schuler, S.C., et al., Extensive remodeling of the extracellular matrix during aging contributes to age-dependent impairments of muscle stem cell functionality. Cell Rep, 2021. 35(10): p. 109223.

40. De Micheli, A.J., et al., Single-cell transcriptomic analysis identifies extensive heterogeneity in the cellular composition of mouse Achilles tendons. Am J Physiol Cell Physiol, 2020. 319(5): p. C885–C894.

41. Lang, F., et al., Dynamic changes in the mouse skeletal muscle proteome during denervation-induced atrophy. Dis Model Mech, 2017. 10(7): p. 881–896.

42. Otten, C., et al., Xirp proteins mark injured skeletal muscle in zebrafish. PLoS One, 2012. 7(2): p. e31041.

43. Molt, S., et al., Aciculin interacts with filamin C and Xin and is essential for myofibril assembly, remodeling and maintenance. J Cell Sci, 2014. 127(Pt 16): p. 3578–92.

44. Leber, Y., et al., Filamin C is a highly dynamic protein associated with fast repair of myofibrillar microdamage. Hum Mol Genet, 2016. 25(13): p. 2776–2788.

45. Tsay, H.J. and J. Schmidt, Skeletal muscle denervation activates acetylcholine receptor genes. J Cell Biol, 1989. 108(4): p. 1523–6.

46. Yumoto, N., S. Wakatsuki, and A. Sehara-Fujisawa, The acetylcholine receptor gamma-to-epsilon switch occurs in individual endplates. Biochem Biophys Res Commun, 2005. 331(4): p. 1522–7.

47. Proietti, D., et al., Activation of skeletal muscle-resident glial cells upon nerve injury. JCI Insight, 2021. 6(7).

48. Love, F.M. and W.J. Thompson, Schwann cells proliferate at rat neuromuscular junctions during development and regeneration. J Neurosci, 1998. 18(22): p. 9376–85.

49. Sugiura, Y. and W. Lin, Neuron-glia interactions: the roles of Schwann cells in neuromuscular synapse formation and function. Biosci Rep, 2011. 31(5): p. 295–302.

50. Huang, X., J. Jiang, and J. Xu, Denervation-Related Neuromuscular Junction Changes: From Degeneration to Regeneration. Front Mol Neurosci, 2021. 14: p. 810919.

51. Min, B.D., et al., Advancing muscle aging and sarcopenia research through spatial transcriptomics. Osteoporos Sarcopenia, 2025. 11(2 Suppl): p. 22-31.

52. Umek, N., et al., In situ spatial transcriptomic analysis of human skeletal muscle using the Xenium platform. Cell Tissue Res, 2025. 399(3): p. 291–302.

53. Hao, Y., et al., Dictionary learning for integrative, multimodal and scalable single-cell analysis. Nat Biotechnol, 2023.

54. Hulsen, T., DeepVenn--a web application for the creation of area-proportional Venn diagrams using the deep learning framework Tensorflow. js. arXiv preprint arXiv:2210.04597, 2022.

55. Si, Y., et al., FICTURE: Scalable segmentation-free analysis of submicron resolution spatial transcriptomics. bioRxiv, 2023.

